# Enhancing Pan-cancer Spatial Transcriptomics at Single-cell Resolution with STPAINTER

**DOI:** 10.64898/2026.02.11.704553

**Authors:** Yuhang Yang, Yiming Luo, Kai Zhang, Zaixi Zhang, Haoxin Peng, Chenlin Cao, Qi Liu, Bin Ma, Yang Chen, Lin Shen, Enhong Chen

## Abstract

Subcellular spatial transcriptomics technologies offer unprecedented views of tissue architecture but are fundamentally constrained by sparse gene panels and limited detection sensitivity. Current computational enhancement strategies typically rely on tissue-matched single-cell RNA sequencing (scRNA-seq) references and necessitate computationally intensive retraining for each dataset, impeding their scalability and clinical applicability. Here, we present STPAINTER, a conditional generative model that leverages a massive pretraining pan-cancer scRNA-seq atlas to universally enhance spatial transcriptomics data. Built upon a latent diffusion architecture with stochastic differential equation-guided generation, STPAINTER learns a universal manifold of cellular states to reconstruct genome-wide expression profiles from sparse spatial measurements. Uniquely, our pretraining paradigm enables zero-shot generalization and empowers downstream tasks by providing imputed transcriptomes and informative latent variables to enhance resolution at both the gene and cluster levels. Applied STPAINTER upon 6 spatial transcriptomics datasets of different cancer types, we demonstrate that our model empowers high-fidelity downstream analyses, including fine-grained subpopulation clustering and pathway enrichment. Furthermore, cross-validation with spatially resolved proteomics (CODEX) confirms the biological veracity of the imputed cellular landscapes. STPAINTER provides a robust, scalable framework for decoding complex tumor microenvironments without the need for auxiliary sequencing data.

## Introduction

The advent of spatial transcriptomics (ST) has enabled the systematic interrogation of gene expression within intact tissues, preserving spatial organization that is lost in dissociative single-cell RNA sequencing (scRNA-seq) workflows^1,2^. While scRNA-seq provides a comprehensive molecular census of cellular identities and states, it lacks spatial context that is essential for understanding tissue architecture and cell–cell interactions. Recent advances in high-resolution ST technologies, particularly image-based platforms such as ×10 Genomics Xenium and NanoString CosMx^3,4^, have pushed spatial profiling to true single-cell and subcellular resolution with single-molecule sensitivity^5^. In contrast to sequencing-based ST methods that measure transcript abundance at multi-cellular spot levels^6–8^, image-based ST enables precise localization of transcripts within individual cells, closely aligning its data structure with that of scRNA-seq^4^. This convergence allows established single-cell analytical frameworks to be extended into the spatial domain, positioning image-based ST as a critical modality for in situ dissection of complex tissue ecosystems such as the tumor microenvironment.

Despite these advances, single-cell resolution ST data suffer from technical limitations that hinder downstream analysis. A primary challenge is the limitations of RNA capture, resulting in high sparsity and technical noise, which obscures subtle biological signals^9,10^. Furthermore, image-based probe hybridization techniques used for single-cell resolution ST often rely on targeted gene panels, capturing only a fraction of the transcriptome up to 5000 genes in Xenium 5k. Consequently, the sparsity and limited coverage compromise the accuracy of unsupervised clustering and annotation, the identification of differentially expressed genes (DEGs) and spatial niche characteristics, particularly for low-abundance transcripts.

To surmount these inherent constraints of ST data, researchers have developed computational strategies for gene imputation, which integrate transcriptomic profiles from scRNA-seq to enhance spatial fidelity. These approaches broadly bifurcate into **alignment-based**^11–17^ and **generative-based**^18–22^ paradigms. The former focuses on establishing direct correspondence between modalities; notable examples include Seurat^11^, which utilizes probabilistic transfer anchors to align reference phenotypes, Tangram^12^, which applies a deep learning framework to optimally map scRNA-seq cells to spatial spots by minimizing feature-level alignment costs, NovoSpaRc^15^, which postulates a structural correspondence between transcriptomic similarity and physical proximity to probabilistically map cells via Gromov-Wasserstein optimal transport, and SpaOTsc^16^, which employs structured and unbalanced optimal transport to address population mismatches, deriving a spatial metric to reconstruct cell locations and infer intercellular communication networks. Conversely, generative-based methods transcend simple mapping by leveraging deep architectures to explicitly learn and internalize intrinsic cellular gene expression patterns. This category encompasses diverse advanced frameworks: gimVI^18^ pioneers the use of variational autoencoders for probabilistic integration, stDiff^21^ harnesses diffusion models to reconstruct spatial gradients, and the recently introduced SpaIM^22^ utilizes style transfer learning to effectively disentangle modality-specific styles from shared biological content. Ultimately, this generative paradigm transcends rigid reference retrieval, synthesizing complex spatial patterns directly from learned data distributions.

Notwithstanding the significant strides achieved by prior gene imputation strategies, critical limitations persist that impede their broad applicability and downstream efficacy. **Firstly**, the majority of existing models operate under a strictly supervised paradigm, necessitating tissue-matched scRNA-seq references. This dependency renders them incapable of generalizing to unseen tissues, compelling computationally expensive retraining for every new biological scenario. **Secondly**, although these methods successfully expand the gene panel, the resulting imputed profiles often retain the high dimensionality, and stochastic noise characteristic of ST data. Consequently, the enhanced data remains suboptimal for direct usage, necessitating secondary dimensionality reduction (e.g., PCA or encoders) to extract usable biological signals for downstream analysis. **Finally**, most legacy approaches are optimized for coarser spot-level resolutions. When applied to cell-level data characterized by higher throughput yet limited known genes, these methods frequently suffer from computational inefficiencies and noise amplification.

In this work, we introduce **STPAINTER**, a latent diffusion model pretrained on massive pan-cancer scRNA-seq datasets to universally enhance ST data across diverse histological landscapes. Leveraging a pretraining atlas, STPAINTER enables direct application to diverse tumors without retraining. Methodologically, STPAINTER employs a conditional latent diffusion framework^23^ to capture shared and distinct gene expression patterns across malignancies. It compresses high-dimensional gene profiles into a compact latent space, where we simulate a Stochastic Differential Equation (SDE)^24^ to enhance features before decoding them back to the gene space. Ultimately, our model yields dual outputs for each cell: a latent embedding for tasks such as fine-celltype clustering, and an imputed gene matrix for differential expression analysis. Furthermore, by performing inference within latent space, our approach achieves superior computational efficiency. Extensive benchmarking across diverse cancer types and platforms confirms STPAINTER’s superiority over existing methods. Subsequent analyses further substantiate its utility for downstream tasks. STPAINTER is available as open-source software on GitHub (https://github.com/yyh030806/stPainter), with detailed but user-friendly tutorials demonstrating its capabilities in enhancing the utility of single-cell resolution ST.

## 2.Results

### 2.1 Overview of STPAINTER

We introduce STPAINTER, a generative framework designed to transcend the boundaries of conventional spatial imputation by establishing a scalable pretrained and inference paradigm (Fig. 1a). Unlike existing methods that necessitate strictly paired references and supervised mapping, thereby severely restricting generalizability, STPAINTER leverages a massive pan-cancer scRNA-seq atlas, encompassing **1.3 million cells** across **21 diverse cancer types**, to learn a universal cellular prior (Fig. 1b). This data-centric approach enables the model to capture a genome-wide manifold of cellular states, facilitating zero-shot generalization to diverse histological landscapes without the need for dataset-specific retraining or matched reference data. By learning from this vast heterogeneity, STPAINTER effectively bridges the gap between the high-resolution ST and the gene coverage of single-cell sequencing.

**Fig. 1.**
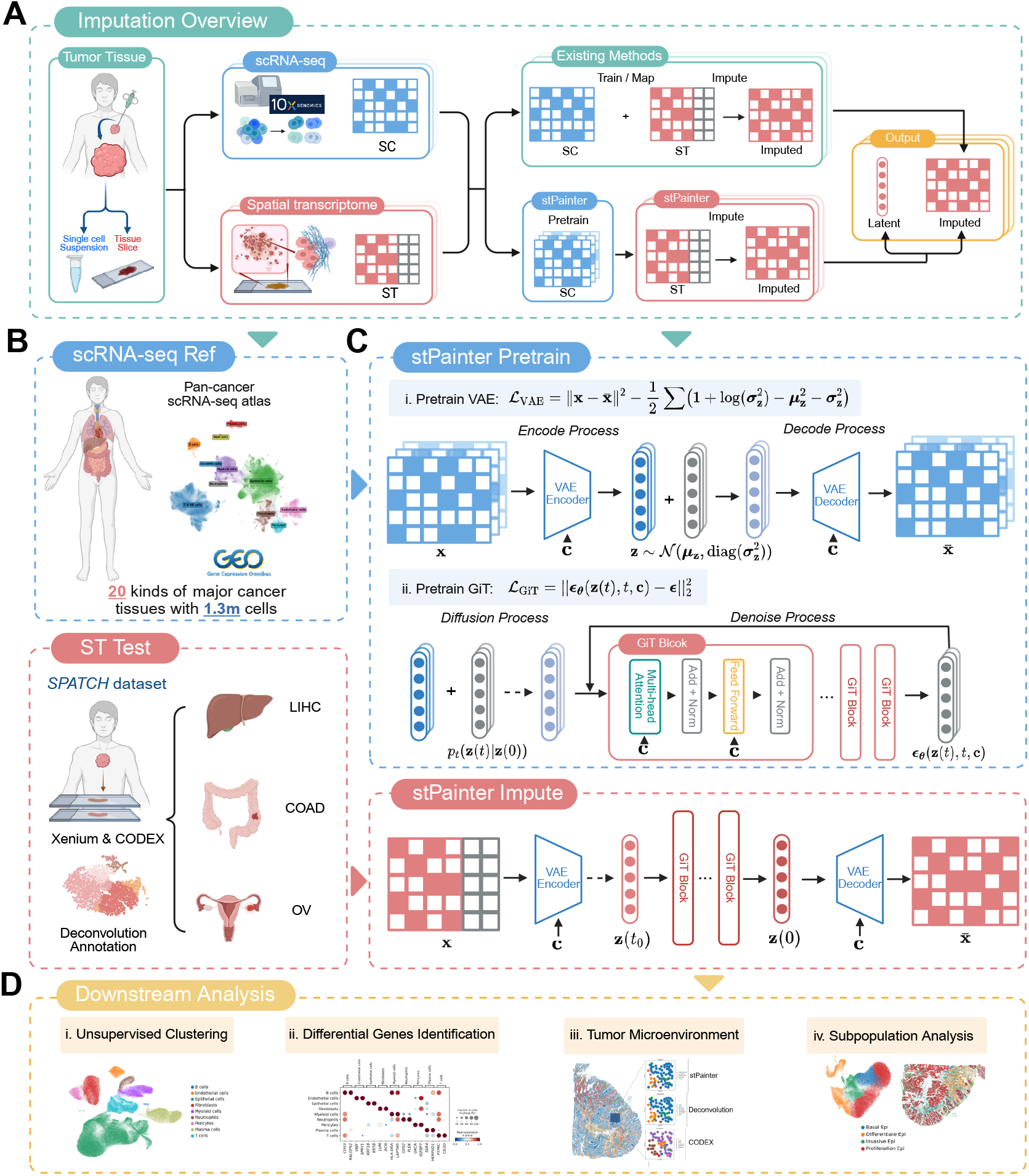
Overview of STPAINTER. **a**, STPAINTER adopts a pretraining paradigm on unpaired scRNA-seq data to generate imputed expression and latent embeddings, unlike supervised methods requiring paired references. **b**, pretrained on a pan-cancer atlas, the model generalizes to ST data across diverse cancer types without retraining. **c**, STPAINTER decouples pretraining and imputation; the training phase sequentially optimizes the VAE and GiT to learn generative priors, enabling the model to be directly applied for inference. **d**, Enhanced data supports robust downstream analyses, including clustering, differential expression, and microenvironment characterization.

To model this complex distribution, STPAINTER employs a latent diffusion architecture (Fig. 1c). First, a Variational Autoen-coder (VAE) maps high-dimensional expression profiles into a compact latent manifold, filtering out technical noise while preserving biological variance. Within this manifold, we train a Gene Diffusion Transformer (GiT) as the conditional generative model, which explicitly conditions the generation process on cancer type. Mathematically, the inference is formulated as a guided generation problem via the SDE. Instead of generating cells from pure noise, we conceptualize the ST measurement as a corrupted state. The model initializes the reverse diffusion process with the latent representation of the observed ST data and iteratively refines it using the learned generative priors. This process effectively projects partial spatial measurements onto the high-fidelity cellular manifold, recovering missing information consistent with the global biological context.

Ultimately, STPAINTER empowers comprehensive downstream characterization through its dual outputs: a denoised latent embedding and a fully imputed genome-wide expression matrix (Fig. 1d). The topology-preserving latent embedding significantly enhances unsupervised clustering and subpopulation analysis, resolving fine-grained cellular heterogeneity often obscured by sparsity. Simultaneously, the imputed high-fidelity matrix enables the precise identification of differentially expressed genes (DEGs), facilitating the discovery of cell-group transcriptomic markers. By recovering complete gene signatures in their native spatial context, STPAINTER further allows for an in-depth dissection of the TME and spatial niches with unprecedented clarity.

### 2.2 STPAINTER enables high-fidelity recovery of spatial gene expression

To evaluate the capability of STPAINTER in recovering spatial gene expression, we implemented two variants of our model: STPAINTER-100 (using a 100 latent size) and STPAINTER-50 (using a 50 latent size). We conducted comparative experiments on three cancer types with paired scRNA-seq references: Colorectal Cancer (COAD), Ovarian Cancer (OV), and Liver Hepatocellular Carcinoma (LIHC). We benchmarked performance against six existing baseline methods: Tangram^12^, NovoSpaRc^15^, SpaOTsc^16^, gimVI^18^, stDiff^21^, and SpaIM^22^. Performance was comprehensively assessed using four quantitative metrics: Pearson Correlation Coefficient (PCC), Structural Similarity Index Measure (SSIM), Root Mean Square Error (RMSE), and Jensen-Shannon divergence (JS). Additionally, to demonstrate the model’s generalizability to scenarios lacking matched references, we validated STPAINTER on three datasets (Breast invasive carcinoma (BRCA), Non-Small Cell Lung Cancer (NSCLC), Prostate Adenocarcinoma (PRAD)) without paired scRNA-seq data, as shown in Fig. S3.

Quantitative benchmarking across varying numbers of highly variable genes (HVGs) further underscores the robust performance and scalability of STPAINTER (Fig. 2a). Specifically, on the COAD dataset, both STPAINTER-100 and STPAINTER-50 consistently surpassed all baseline methods across the four evaluation metrics. As the number of imputed genes scaled from 10 to 300, STPAINTER-100 maintained a superior Pearson Correlation PCC. This metric peaked at 0.31 for the top 100 genes, which represents a significant margin over the leading generative baseline gimVI (PCC = 0.24) and alignment-based Tangram (PCC = 0.24). The advantage of STPAINTER was most pronounced in structural fidelity as measured by the SSIM. While STPAINTER variants sustained SSIM scores above 0.30 across the entire gene range, other methods, particularly stDiff and SpaIM, yielded values below 0.01. These near-zero results indicate an inability to reconstruct coherent spatial expression manifolds. Furthermore, STPAINTER achieved the lowest global error (RMSE = 1.16) and statistical divergence (JS = 0.48) at the 100 HVG scale. These results suggest that the imputed profiles closely mirror the absolute abundance and distribution of the ground-truth transcripts. Notably, the performance of STPAINTER remained remarkably stable as the complexity of the gene panel increased, whereas baseline methods often exhibited diminishing returns. This consistency highlights the effectiveness of the universal cellular prior in regularizing the imputation process, allowing STPAINTER to generalize high-fidelity recovery across varying genomic scales.

**Fig. 2.**
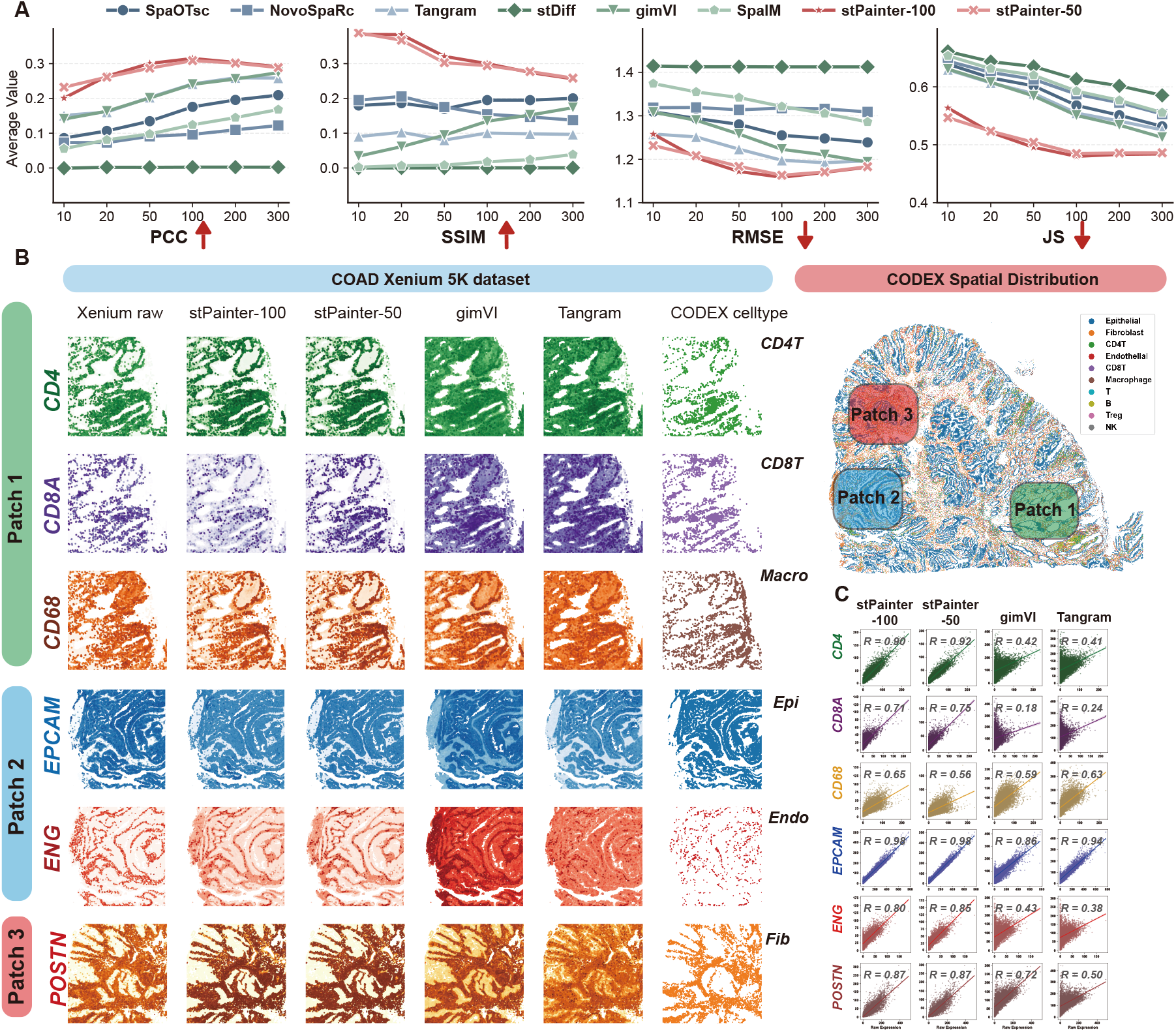
High-fidelity recovery of spatial gene expression by STPAINTER. **a**, Quantitative benchmarking of gene expression recovery on the COAD dataset. Performance was evaluated using four metrics: Pearson Correlation Coefficient (PCC), Structural Similarity Index Measure (SSIM), Root Mean Square Error (RMSE), and Jensen-Shannon divergence (JS). **b**, Visual comparison of spatially resolved marker gene expression across three representative tissue patches. **c**, Gene-wise scatter plots showing the correlation between imputed expression levels and raw Xenium measurements for selected marker genes. R-value indicates the PCC calculated within 100 *µ*m × 100 *µ*m spatial regions.

We further examined the imputation quality of biologically significant marker genes to validate spatial fidelity. As illustrated in Fig. 2b, STPAINTER accurately reconstructed the spatial distributions of key immune and structural markers, including *CD4* and *CD8A* (T cells), *CD68* (macrophages), *EPCAM* (epithelial cells), *ENG* (endothelial cells), and *POSTN* (fibroblasts). The imputed gene expression patterns generated by STPAINTER showed high concordance with both the ground-truth Xenium measurements and the cell-type spatial distributions derived from CODEX protein imaging. To further assess whether the imputed marker genes faithfully reflected the in situ tumor microenvironment, we quantified the correlation of key gene expression profiles within corresponding 100 *µ*m ×100 *µ*m spatial grids. As shown in Fig. 2c, STPAINTER achieved the highest correlation with the ground-truth data, particularly for T-cell markers *CD4* and *CD8A*, as well as the vascular marker *ENG*. In contrast, baseline methods frequently produced blurred spatial signals or artifacts, which compromised the precise localization of these biologically relevant markers. Finally, we extended this evaluation to the OV and LIHC datasets to ensure robustness across different tissue types. Consistent with the findings in COAD, STPAINTER demonstrated robust performance advantages in these cancer types as well, the detailed comparative results of which are presented in Fig. S2a-d.

### 2.3 STPAINTER enhances cell-type identification and spatial mapping

Accurate cell type identification is a prerequisite for interpreting ST data. We evaluated whether the latent representations and imputed profiles generated by STPAINTER could improve unsupervised clustering performance compared to raw data and baseline methods.

To quantitatively evaluate the capacity of STPAINTER in resolving spatial domains, we benchmarked its two variants (STPAINTER-100 and STPAINTER-50) against six existing methods using four complementary clustering metrics: Adjusted Rand Index (ARI), Adjusted Mutual Information (AMI), Homogeneity (Homo), and Normalized Mutual Information (NMI). The quantitative results on the COAD dataset (Fig. 3a) demonstrate that STPAINTER consistently outperforms all baseline methods across all four metrics. Specifically, STPAINTER-100 achieved the highest scores, with an ARI of 0.85, AMI of 0.70, Homo of 0.63, and NMI of 0.70. Even with a reduced latent dimension, STPAINTER-50 maintained competitive performance (ARI = 0.80, AMI = 0.65, Homo = 0.55, NMI = 0.65), still significantly exceeding the best-performing generative baseline, SpaIM (ARI = 0.57, AMI = 0.52, Homo = 0.37, NMI = 0.52), and alignment-based methods like Tangram (ARI = 0.56, AMI = 0.50, Homo = 0.35, NMI = 0.50). In contrast, other baseline tools such as gimVI and stDiff showed substantially lower scores, while NovoSpaRc and SpaOTsc failed to provide biologically meaningful clustering results, yielding near-zero values across the board. These consistent gains across multiple metrics suggest that by leveraging a universal cellular prior, STPAINTER effectively reconstructs a high-fidelity manifold that is essential for accurate and robust tissue segmentation in complex cancer landscapes.

**Fig. 3.**
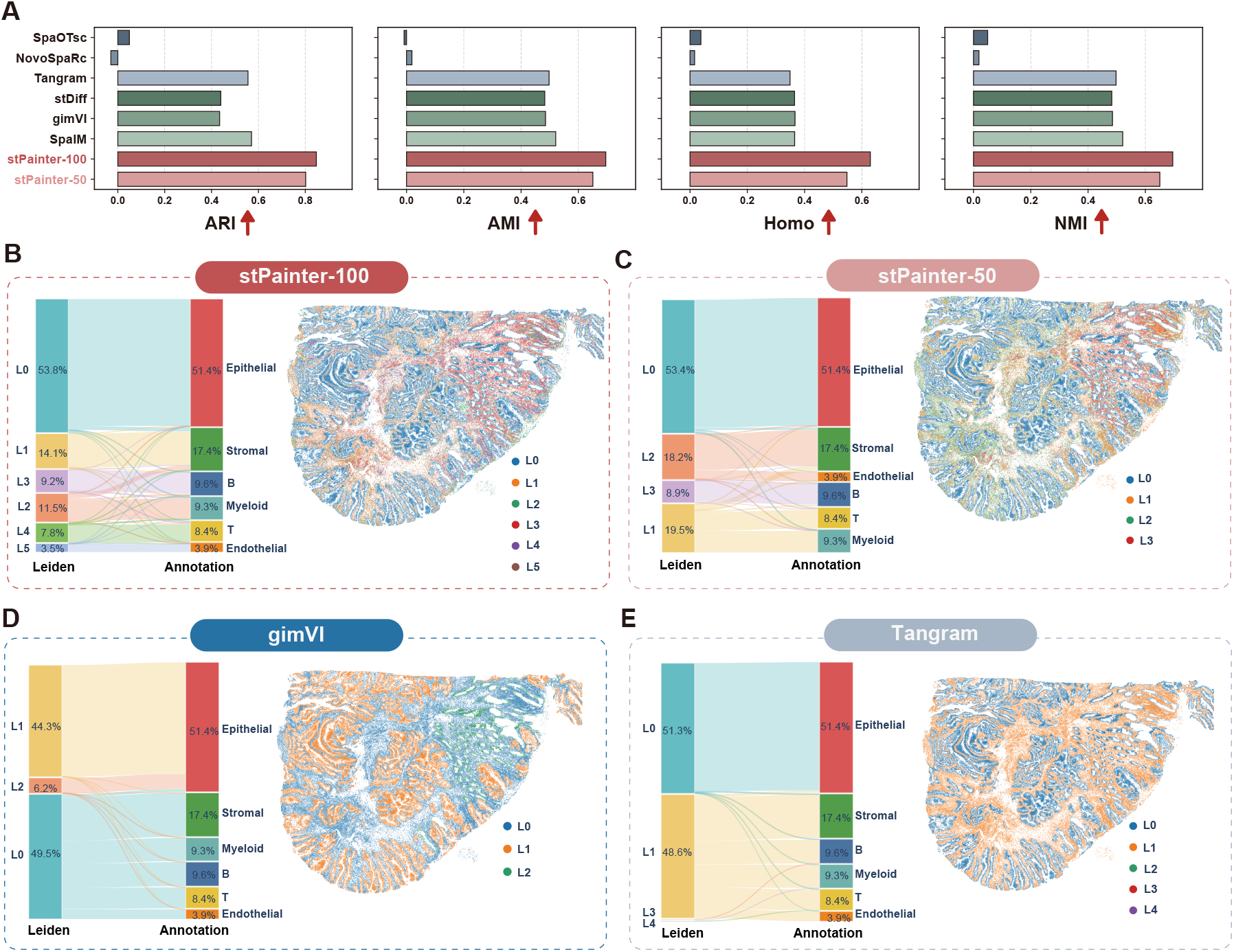
Benchmarking of unsupervised clustering with imputed xenium data. **a**, Quantitative evaluation of clustering performance on the COAD Xenium dataset across four metrics: Adjusted Rand Index (ARI), Adjusted Mutual Information (AMI), Homogeneity (Homo), and Normalized Mutual Information (NMI). **b–e**, Sankey diagrams and corresponding spatial cluster maps illustrating the mapping between unsupervised Leiden clusters (left nodes) and transfer-annotation cell types (right nodes). **b, c**, Clustering results derived from STPAINTER-100 (**b**) and STPAINTER-50 (**c**) latent embeddings reveal clean separation of cell populations and spatially coherent tissue structures. **d, e**, In contrast, baseline methods such as gimVI (**d**) and Tangram (**e**) exhibit higher entanglement between clusters and fragmented spatial domains, indicating reduced biological specificity.

To assess the concordance between unsupervised clusters and biological identities, we used Sankey diagrams to map Leiden clusters derived from the imputed data to transfer-annotation labels (Fig. 3b–e). At low resolution (r = 0.02), clustering results from STPAINTER-100 and STPAINTER-50 showed clear and coherent mappings for major cell types (Fig. 3b, c). In contrast, under the same resolution, baseline methods such as gimVI and Tangram produced more fragmented clustering patterns.

Their Sankey diagrams exhibited pronounced entanglement, with individual cell types split across multiple noisy clusters or, conversely, distinct cell types merged into a single cluster (Fig. 3d, e).

### 2.4 STPAINTER resolves spatial cellular heterogeneity from imputed transcriptomes

Single-cell resolution ST platforms, such as Xenium 5k, pose substantial challenges for downstream analyses, including unsupervised clustering and cell type annotation^10^. We show that gene imputation with STPAINTER enables robust and biologically coherent downstream analysis. Applying STPAINTER-100 to Xenium COAD datasets, we reconstructed genome-wide transcriptomes comprising 10,000 genes together with compact latent representations (Fig. 4a). Latent-driven UMAP embeddings revealed well-separated and structured cell clusters that closely resemble those obtained from high-quality scRNA-seq data. These clusters exhibited strong concordance with cell type labels derived from transfer annotation. Moreover, the selective and robust expression of canonical marker genes, such as *KRT19* in epithelial cells and *PDGFRB* in smooth muscle cells, further confirms that STPAINTER enhances transcriptomic resolution while preserving biological fidelity.

**Fig. 4.**
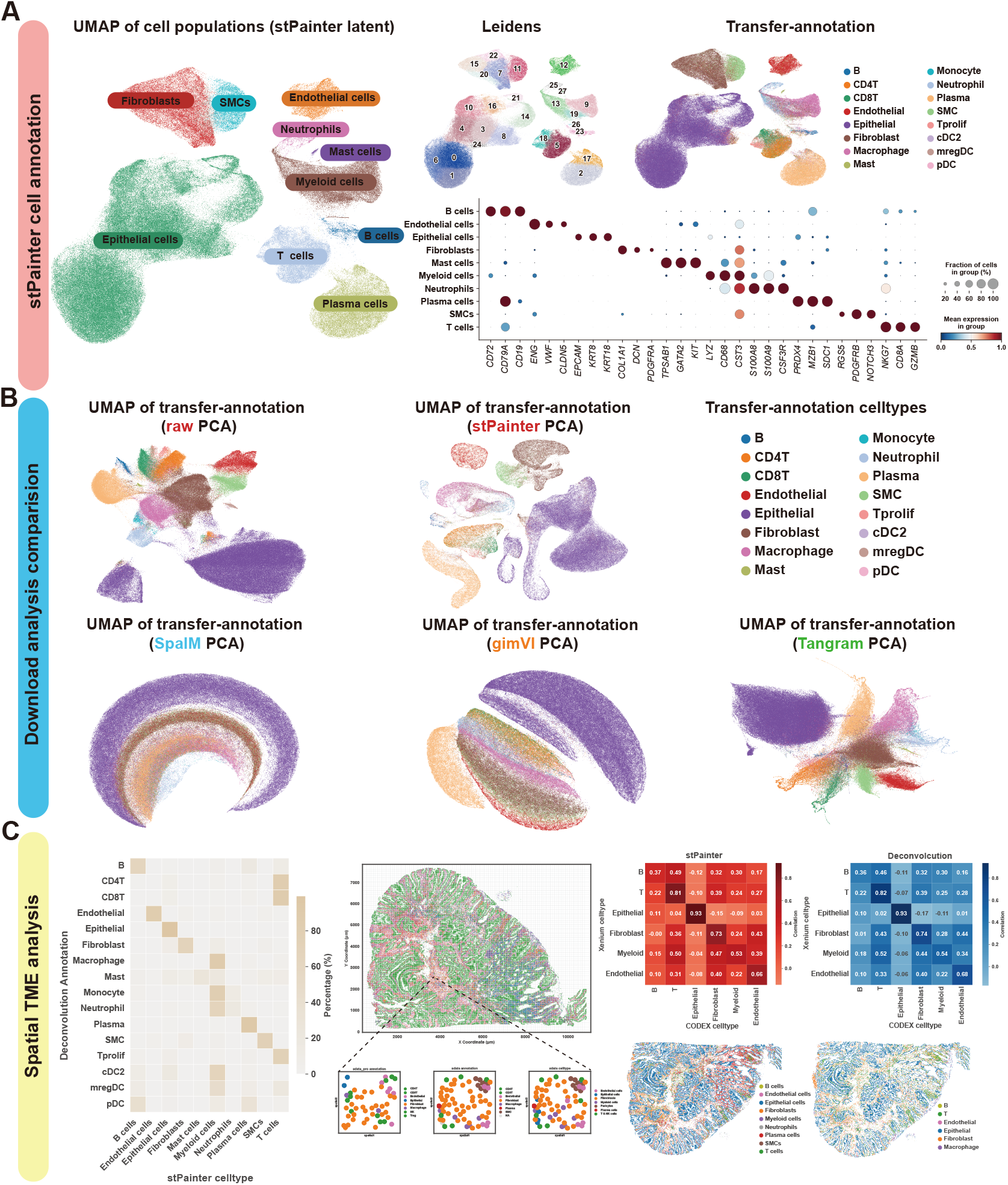
Resolution of spatial cellular heterogeneity and TME architecture. **a**, UMAP projection of cell populations derived from STPAINTER latent embeddings and corresponding dot plot of lineage-defining markers. **b**, Comparative UMAP topologies generated by different computational workflows (STPAINTER-100, SpaIM, gimVI, and Tangram). **c**, Spatial coherence and lineage verification. The heatmap of cell-type proportions and correlation matrices (Pearson’s *r* index) validate the concordance between STPAINTER-imputed cellular compositions and ground-truth CODEX proteomics across tissue patches.

We further benchmarked STPAINTER against baseline tools and the standard Xenium processing workflow (PCA-based clustering utilizing transfer-annotation labels)^25^. In contrast to the raw data and other imputation methods, only STPAINTER recovered UMAP topologies rich in biological interpretability (Fig. 4b). Notably, competing methods frequently exhibited over-smoothed trajectories in UMAP space. These structures likely represent technical artifacts resulting from the generation of dense matrices without adequately modeling the inherent sparsity of ST data, rather than true biological gradients. Consequently, these artifacts obscured distinct cell populations, leading to inferior clustering performance compared to STPAINTER.

Finally, we validated the spatial fidelity of the imputed data within the tissue microenvironment. We found that cell type annotations derived from STPAINTER exhibited high consistency with transfer-annotation results. To quantitatively assess this, we segmented the tissue sections into 100 *µ*m ×100 *µ*m patches and compared the cellular composition against ground truth data from CODEX imaging of adjacent serial sections. Our analysis revealed that STPAINTER-derived annotations achieved a high correlation with CODEX data, comparable to transfer-annotation. This high degree of concordance was particularly evident in key lineages, including epithelial cells and T cells, validating the accuracy of STPAINTER in resolving spatial cellular distributions.

### 2.5 STPAINTER enables fine-grained dissection of cell subpopulations

Fine-grained cell annotation is pivotal for dissecting tissue heterogeneity in solid tumors, particularly within the immune and epithelial compartments^26^. However, identifying rare subpopulations in raw ST data remains challenging, where lineage-specific marker genes often suffer from low detection efficiency^27^. To overcome this limitation, we isolated T cells, myeloid cells, and epithelial cells from the STPAINTER-imputed COAD Xenium dataset and performed high-resolution sub-clustering with STPAINTER-latent.

Within the T cell compartment, we resolved distinct subtypes, including CD4^+^ memory T cells (Tm), naive T cells (Tn), regulatory T cells (Treg), CD8^+^ effector T cells (Teff), exhausted T cells (Tex), and proliferating T cells (Tprolif) (Fig. 5a). STPAINTER significantly enhanced the expression of canonical markers defining these lineages—such as *FOXP3* in Tregs, *CCR7* in CD4^+^ Tn, and *CXCR5* in CD8^+^ Tex—thereby recovering unique biological identities that were otherwise obscured^28^. To validate spatial fidelity, we aligned these annotations with protein-based T cell clusters (CD4^+^ T, CD8^+^ T, Treg, and others) derived from adjacent CODEX sections. Pearson correlation analysis of cell densities within 100 *µ*m ×100 *µ*m patches revealed a high degree of concordance between the STPAINTER-derived subpopulations and the ground-truth protein imaging, confirming that our model accurately recapitulates the spatial architecture of the TME (Fig. 5b,d). Similarly, fine-grained resolution was achieved in the myeloid compartment. Beyond major populations such as macrophages, monocytes, neutrophils, and mast cells, the model successfully recovered rare subsets like regulatory dendritic cells (mregDC), characterized by the specific expression of *CCR7* (Fig. 5e-h)^29^.

**Fig. 5.**
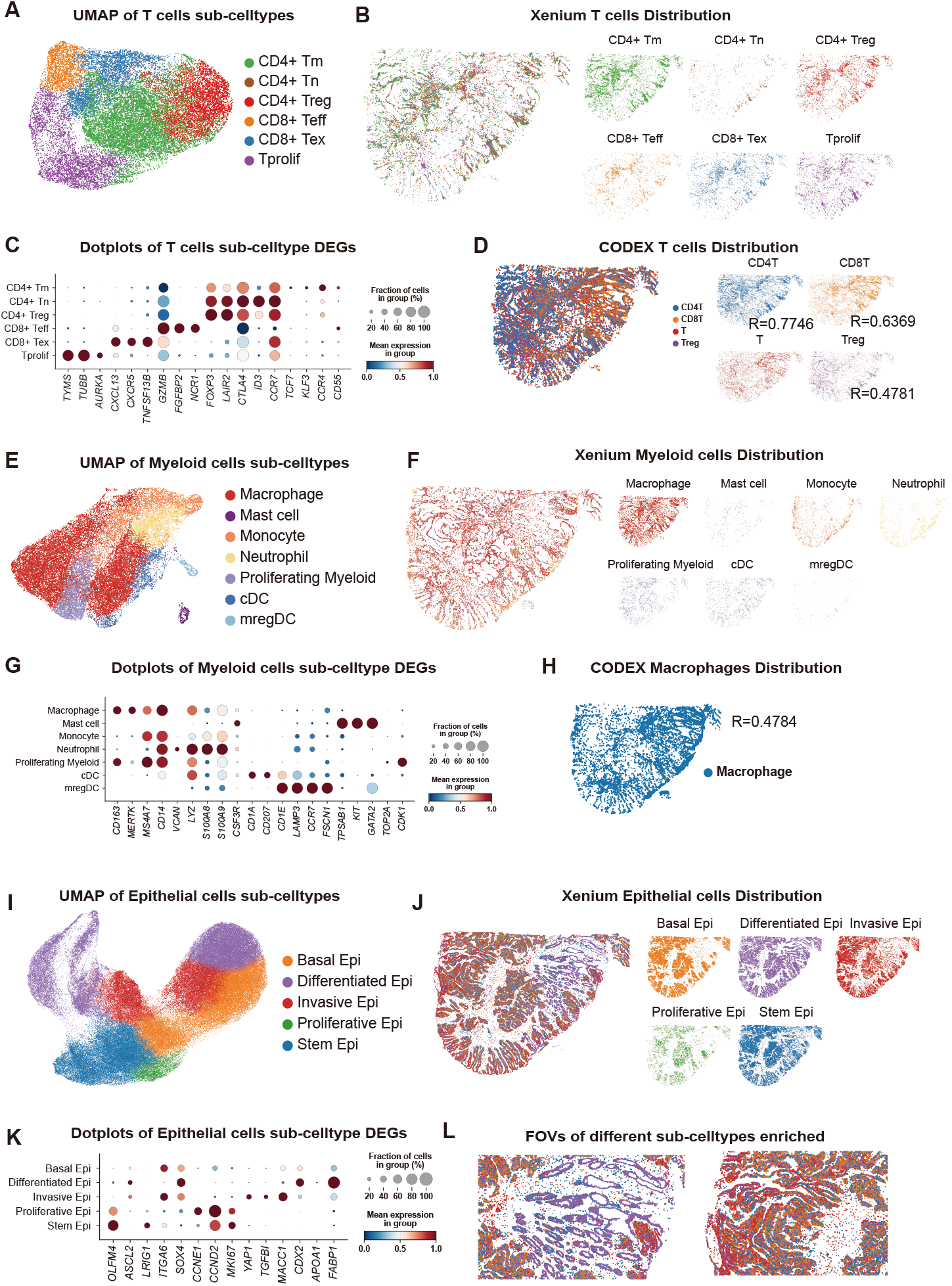
Multi-modal characterization of fine-grained cell sub-lineages and their spatial niches. **a–d**, T cell lineage dissection via UMAP visualization (**a**), Xenium-based spatial mapping (**b**), dot plot of T cell DEGs (**c**), and CODEX proteomic validation (**d**). **e–h**, Myeloid cell lineage dissection via UMAP visualization (**e**), Xenium-based spatial mapping (**f**), dot plot of myeloid cell DEGs (**g**), and CODEX proteomic validation (**h**). **i–l**, Epithelial cell lineage dissection via UMAP visualization (**i**), Xenium-based spatial mapping (**j**), dot plot of epithelial cell DEGs (**k**), and representative spatial enrichment within specific fields-of-view (FOVs) (**l**).

Epithelial cells are fundamental to understanding tumor heterogeneity. We stratified epithelial cells into five distinct functional states: Differentiated, Basal, Stem, Proliferative, and Invasive Epithelial cells (Fig. 5i). Differential expression analysis confirmed lineage specificity: Differentiated cells expressed *CDX2*^30^, Basal cells were marked by *ITGA6*^31^, while Stem populations showed elevated *OLFM4*^32^. Notably, the Invasive cluster was enriched for genes associated with Epithelial-Mesenchymal Transition (EMT), including *TGFB1* and *MACC1* (Fig. 5k). Spatial mapping revealed distinct morphological contexts: Differentiated cells formed organized glandular structures, whereas Invasive cells were preferentially localized at the invasive tumor front (Fig. 5j, l)^33^. These findings demonstrate that STPAINTER-derived annotations are biologically coherent and accurately reflect the functional landscape of tumor progression.

Finally, we evaluated the generalizability of STPAINTER-100 across an expanded spectrum of malignancies by extending our analyses to the remaining SPATCH datasets (OV and LIHC) and multiple independent public Xenium datasets, including PRAD, NSCLC, and CESC (Fig. S7–S10). In all cases, STPAINTER facilitated robust downstream workflows, achieving a level of resolution in major and fine-grained subpopulation identification comparable to that observed in COAD. These results underscore the robustness of STPAINTER, validating its capacity to universally enhance ST data and streamline analysis across diverse cancer histologies.

## 3 Discussion

High-resolution ST^25^ is fundamentally constrained by stochastic RNA capture and restricted gene panels, necessitating the integration of whole-transcriptome scRNA-seq to restore biological signals and enhance data fidelity. However, existing imputation frameworks^12,18^ are hampered by a sample-specific supervised paradigm that requires tissue-matched references and repetitive retraining for every new experiment, a dependency that severely restricts their scalability and clinical applicability. Here, we developed STPAINTER, a conditional latent diffusion model that shifts the field toward a pretrained paradigm. By leveraging a massive pan-cancer atlas of 1.3 million cells across 21 diverse malignancies, STPAINTER enables high-fidelity, zero-shot ST data enhancement without the need for auxiliary matched data or retraining, providing a convenient and effective (Fig. S5) tool for ST data analysis.

The technical efficacy of STPAINTER is grounded in its conditional latent diffusion architecture, which formulates gene expression enhancement as a guided traversal on a learned cellular manifold. By encoding cancer-type metadata as a conditional prior (*c*), the model constrains the generative dynamics to a biologically plausible subspace of the global cellular state space. In contrast to conventional approaches that operate directly in the high-dimensional and noise-contaminated gene expression space, STPAINTER performs SDE-guided generation within a highly compressed, denoised latent manifold (*z*)^24^. This procedure iteratively maps sparse and corrupted spatial measurements back onto the high-fidelity distribution learned from a pretrained pan-cancer atlas(Fig. S5) The latent representations produced by STPAINTER constitute a topology-preserving embedding that retains the principal determinants of cellular identity while attenuating technical artifacts. By compressing complex transcriptomic variability into a low-dimensional latent embedding (e.g., 50 or 100 dimensions), the model establishes a stable basis for unsupervised manifold learning and high-resolution cluster delineation, which would otherwise be hindered by the sparsity and noise of raw measurements^34^. In parallel, the generative decoder reconstructs a continuous, genome-wide expression matrix, thereby enabling comprehensive downstream transcriptomic analyses. This dual-layer enhancement strategy couples compact latent embeddings with high-dimensional probabilistic imputations, allowing STPAINTER to jointly optimize macroscopic tissue-level segmentation and microscopic gene-level reconstruction.

Single-cell resolution ST provides gene expression profiles analogous to scRNA-seq, yet critical differences in data characteristics necessitate analytical approaches^25^. Unlike scRNA-seq, where enzymatic dissociation can disproportionately impact fragile cell types like epithelial cells^10,35,36^, xenium technologies often exhibit higher dropout rates for less abundant populations, such as immune cell genes (*CD8A, CD14*) (Fig. S6). Furthermore, signal leakage from abundant neighboring cells can mask the transcriptomic signatures of rarer cell types, complicating their accurate identification^10^. STPAINTER is designed to address these fundamental data quality issues. By enhancing the raw measurements, it expands the gene panel to a comprehensive set of nearly 10,000 genes and mitigates the inherent data sparsity, yielding scRNA-seq-like data structure primed for downstream analysis.

The high levels of noise and sparsity in xenium data also severely compromise downstream computational tasks, particularly the clustering and identification of fine-grained cell subtypes. Conventional methods like PCA often fail to effectively separate biological signals from technical noise, resulting in poor cluster resolution^10^. While many imputation tools exist, they are often not optimized for clustering tasks and can fail to remove technical artifacts^21^. STPAINTER provides a more powerful solution. We demonstrate that performing clustering directly on the latent representations learned by our model, rather than on raw or imputed gene expression, improves both the accuracy and efficiency of cell type identification. The resulting 50-to 100-dimensional latent space robustly captures the salient biological features that define cellular identity. This enables the resolution of fine-grained subtypes with clusters that show strong concordance with known marker genes, and obviates the dependence on a matched scRNA-seq reference for cell annotation.

Despite the robust performance and universal applicability demonstrated by STPAINTER, several avenues remain for further optimization and exploration. Currently, our framework addresses data sparsity based on patterns derived from scRNA-seq; future iterations could explore more adaptive and learnable strategies for sparsity processing that dynamically account for sample-specific technical biases and varying capture efficiencies. Furthermore, while the latent representations produced by our model serve as a high-fidelity basis for downstream clustering, the biological interpretability of these latent dimensions warrants deeper investigation. Uncovering the specific gene-program alignments within this manifold could provide novel insights into the regulatory logic of the tumor microenvironment. Finally, the current pan-cancer atlas faces an inherent challenge of compositional imbalance, with rare malignancies being underrepresented compared to common cancer types. Integrating broader, more balanced multi-omic datasets may further refine STPAINTER ‘s zero-shot generalization and enhance its capacity to resolve the intricate biological granularity essential for downstream mechanistic interrogation.

## 4 Methods

### 4.1 Problem Formulation

Formally, we represent a single-cell resolution transcriptomic dataset as a tuple *𝒥* = (**X, C**), where **X**∈ ℝ^*N*×*G*^ denotes the gene expression matrix for *N* cells and *G* genes, and **C** ∈ ℝ^*N*×*K*^ constitutes the one-hot encoding for *K* distinct cancer types. Accordingly, the *i*-th cell is characterized by the pair (**x**_*i*_, **c**_*i*_). We aim to capture the heterogeneous gene expression patterns intrinsic to different cancer types by pretraining our latent diffusion model STPAINTER on a comprehensive pan-cancer scRNA-seq atlas reference *𝒥*_*sc*_ = (**X**_*sc*_, **C**_*sc*_), which spans a genome-wide feature space *G*_*sc*_. Conversely, we define a target single-cell resolution spatial transcriptomic dataset as *𝒥*_*st*_ = (**X**_*st*_, **C**_*st*_), typically restricted to a specific cancer type with a limited gene panel such that *G*_*st*_ ≪ *G*_*sc*_. Our primary objective is to leverage the learned generative priors to transform the raw *𝒥*_*st*_ into an enhanced 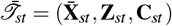. This output comprises the spatially informed expression matrix 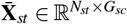 imputed to the full reference gene space, and a compact latent embedding 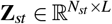 that encapsulates cellular manifolds.

### 4.2 The Architecture of STPAINTER

The architecture of STPAINTER is composed of two synergistic components: a VAE for latent space embedding and a GiT for conditional generative modeling. The VAE is first trained to compress high-dimensional transcriptomic profiles into a compact, structured latent manifold that captures essential biological variance. Subsequently, the GiT functions as our core generative module by operating within this learned manifold. It is designed as a conditional diffusion model to learn the complex probability distribution of cellular states based on auxiliary information, such as cancer type. This decoupled, two-stage approach allows STPAINTER to efficiently handle genome-scale data while leveraging the power of Transformer models for precise, context-aware generation.

#### The Architecture of VAE

Our VAE is inspired by the probabilistic framework of scVI^34^ to learn a latent representation **z** while disentangling it from technical nuisance factors. It consists of an encoder-decoder architecture:

- The encoder network maps the raw gene expression profile **x** and its cancer type **c** to the parameters of the two factorized latent distributions via distinct output heads:

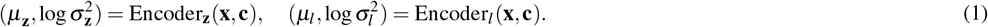

These parameters define the distributions for the biological latent vector **z** ∼ *𝒩* (*µ*_**z**_, diag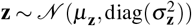) and the scalar library size factor *l* ∼ LogNormal 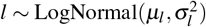. The latent variables are then sampled using the reparameterization trick.

- The decoder network maps the latent state back to the parameters of the Zero-Inflated Negative Binomial (ZINB) distribution via two distinct output heads:

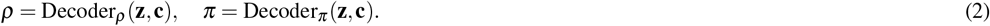

These outputs define the normalized gene frequencies *ρ* and the dropout probabilities *π*. The reconstructed gene count 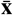 is then drawn from the distribution ZINB(*µ, θ, π*), where the mean is given by *µ* = *l. ρ* and the dispersion *θ* is a globally learnable parameter.

The hyperparameters of the VAE architecture are detailed in Table S2.

#### The Architecture of GiT

The GiT is a conditional diffusion model based on the Diffusion Transformer (DiT)^37^. We specifically adapted its architecture to process 1D latent transcriptomic embeddings, enabling it to model intricate dependencies within the VAE-learned cellular manifold:

- The input to the GiT is the noisy latent vector **z**(*t*). We implement a feature-wise tokenization strategy by treating the latent vector as a sequence of *L* tokens. Each one is projected into a hidden dimension *D* via a learnable linear embedding. Learnable 1D positional embeddings are added to preserve the topological semantics of the latent dimensions, yielding the initial sequence **H**_0_ ∈ ℝ ^*L*×*D*^.
- The core network comprises a stack of Transformer blocks that iteratively denoise the latent representation. We employ an adaptive layer norm zero (adaLN-Zero) mechanism to rigorously condition the process on the cancer type **c** and timestep *t*. For each block, a shared MLP processes the fused condition vector to predict modulation parameters (*γ, β, α*) that are applied to the Multi-Head Self-Attention (MSA) and Feed-Forward Network (FFN) modules:

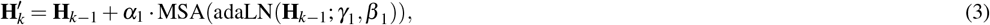

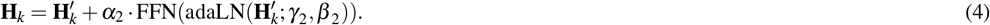

The regression MLP is initialized to zero, rendering each block an identity function at the start of training to stabilize learning dynamics.
- After processing through *n* blocks, the output sequence **H**_*n*_ is projected back to the latent space dimension. The tokens undergo a final adaLN normalization, followed by a linear projection back to the scalar dimension. The sequence is then reshaped to yield the predicted noise required for the diffusion process, formally denoted as:

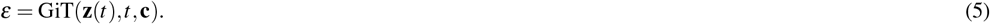

The hyperparameters of the GiT architecture are detailed in Table S3.

### 4.3 Training and Inferencing of STPAINTER

The application of STPAINTER involves two phases: a training phase to learn generative priors from the pan-cancer atlas, and an inferencing phase to enhance target ST data. The training is sequential: we first train the VAE to create a robust latent representation of cellular states. Then, holding the VAE fixed, we train the GiT to model the conditional distribution of these latent states. For inferencing, we employ a guided generation process based on score-based diffusion, which iteratively refines an initial noisy estimate of the target cell’s latent state to be consistent with both the learned priors and the original measurements.

#### Generative Modeling with SDE

The generative process of STPAINTER is formulated through the lens of score-based generative modeling via SDEs^24^. This framework conceptualizes generation as a time-reversed diffusion process. A forward process, defined by a prespecified SDE, systematically perturbs data samples **z**(1) from the complex data distribution *p*_1_ towards a simple prior distribution *p*_0_ (e.g., an isotropic Gaussian) at *t* = 0 over a continuous time interval *t* ∈ [0, 1]. A standard choice for this forward SDE is the Variance Preserving (VP) SDE:

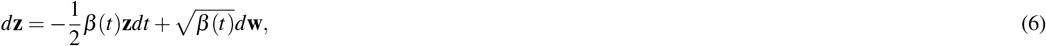

where *β* (*t*) is a positive, monotonically increasing variance schedule defined on *t* ∈ [0, 1], and **w** is a standard Wiener process.

The creative potential lies in the reverse process. The trajectory of this diffusion can be reversed by solving a corresponding reverse-time SDE^38^:

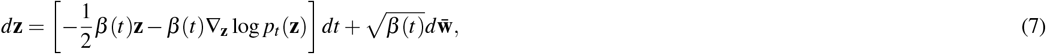

where 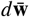 is a Wiener process running backwards in time, and ∇_**z**_ log *p*_*t*_(**z**) is the score function of the perturbed data distribution *p*_*t*_ at time *t*. Simulating this SDE from a noise sample **z**(0) ∼*p*_0_ to *t* = 1 effectively generates a novel sample from the original data distribution *p*_1_. The central challenge is thus to accurately estimate the time-dependent score function, which we accomplish by training our GiT.

#### Training of STPAINTER

The training of STPAINTER is conducted in two sequential stages, ensuring a stable latent manifold is learned before the generative priors are established.

- The VAE is first trained on the pan-cancer scRNA-seq atlas *𝒥*_*sc*_ to learn a compressed and robust latent representation of single-cell transcriptomes. The objective is to maximize the evidence lower bound (ELBO) of the data log-likelihood, which balances reconstruction fidelity with regularization of the latent space:

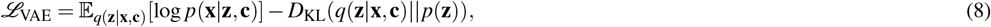

where the first term is the ZINB-based reconstruction loss and the second is the Kullback-Leibler divergence that regularizes the posterior *q* towards a standard normal prior *p*(**z**). Upon convergence, the VAE encoder provides a high-quality mapping from gene expression space to the latent manifold.
- Following VAE convergence, its encoder is used to project the entire pan-cancer atlas into the latent space, creating a dataset of latent vectors **z**_*sc*_. The GiT, parameterized by *θ* and denoted as *ε*_*θ*_, is then trained on this latent dataset. Its objective is to predict the initial noise *ε* added to a latent vector **z**(0) to produce its noisy version **z**(*t*), conditioned on the cancer type **c** and timestep *t*. This is equivalent to learning the score function and is achieved by minimizing the following objective:

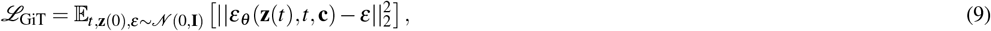

where **z**(*t*) is sampled from the forward process perturbation kernel *p*_*t*_(**z**(*t*)|**z**(0)) and *t* is sampled uniformly from [0, 1].

The training hyperparameters are detailed in Table. S4.

#### Inferencing with STPAINTER

During inferencing with STPAINTER, we employ a guided generative process referenced to Stochastic Differential Editing (SDEdit)^**?**^. This process ensures that the generated profiles are both biologically realistic and faithful to the original, partially measured data. The procedure unfolds in four steps:

- First, we obtain a latent representation **z**_*st*_ of the input limited-gene panel data **x**_*st*_ by passing it through the frozen, pretrained VAE encoder. This maps the low-dimensional measurement onto the learned cellular manifold.
- Instead of initializing the generative process from pure noise (i.e., at *t* = 1), we add a controlled amount of noise to the initial latent state **z**_*st*_. This is achieved by simulating the forward SDE (Eq. 6) starting from **z**(0) = **z**_*st*_ for a specified duration, up to a time *t*_0_ ∈ (0, 1]. This yields a perturbed latent vector **z**(*t*_0_). The hyperparameter *t*_0_ critically governs the trade-off between faithfulness to the original measured expression (lower *t*_0_) and the realism of the imputed profile (higher *t*_0_).
- Starting from the perturbed latent vector **z**(*t*_0_), we then numerically solve the reverse-time SDE (Eq. 7) from *t* = *t*_0_ down to *t* = 0. This is typically performed using a predictor-corrector sampler or a simpler Euler-Maruyama scheme, where each step is guided by the score estimated by our trained GiT, *ε*_*θ*_. This step effectively projects the noisy, partial information onto the manifold of complete, realistic cellular states learned from the pan-cancer atlas.
- The final, denoised latent vector **z**(0) represents the imputed cellular state that optimally balances prior knowledge and measured data. It is then passed through the frozen VAE decoder to deterministically generate the high-dimensional, genome-wide expression profile 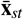, completing the inferencing process.

The sampling hyperparameters are detailed in Table. S5.

### 4.4 Pan-cancer scRNA-seq atlas processing

We aggregated publicly available scRNA-seq datasets encompassing 21 cancer types, including both primary and metastatic tumors. To ensure broad genomic representation across cancer studies, only genes present in all datasets were retained for integration^39^. Cells were excluded if they exhibited any of the following: mitochondrial gene fraction >20%, ribosomal gene fraction >40%, fewer than 300 detected genes, or total UMI counts <500. Highly variable genes were identified using the *Scanpy* implementation of the normalized dispersion method; the top 10,000 most variable genes were selected as targets for downstream imputation. To evaluate the biological fidelity of the integrated atlas, we applied scVI for batch-corrected integration after quality control^34^. The resulting embedding resolved canonical cell subtypes, confirming the atlas’s suitability for pan-cancer analyses.

### 4.5 The down-stream analysis of ST dataset

To facilitate robust downstream analysis, we first performed quality control by excluding the bottom 15% of cells based on total UMI counts. Gene expression imputation was then carried out using STPAINTER. Subsequent analyses followed the standard *Scanpy* workflow: after log-normalization, low-dimensional embeddings were derived using both STPAINTER ‘s latent representation and PCA. Neighborhood graphs were constructed, followed by UMAP dimensionality reduction and Leiden clustering. Given that cell-type annotations were initially assigned using transfer-annotation approach, we assessed the concordance between these transfer-annotation labels and the Leiden clusters. To refine annotation resolution, we performed subclustering and re-annotation of major cell populations and validated their identities by comparing marker gene expression profiles with those from reference single-cell transcriptomic datasets of the same tissue type. Major cell populations were annotated based on the expression of canonical marker genes: *CD8A* (T cells), *CD19* (B cells), *MZB1* (plasma cells), *CD68* (myeloid cells), *FCGR3B* (neutrophils), *PLVAP* (endothelial cells), *COL1A1* (fibroblasts), and *PDGFRB* (SMCs).

To evaluate the impact of STPAINTER imputation on the spatial architecture of the TME, we conducted a spatial coherence analysis based on cell-type annotations. Specifically, we partitioned each tissue section into 100µm×100µm patches and quantified the cellular composition within each patch. Leveraging spatially resolved CODEX proteomics data from adjacent sections as a ground-truth reference, we computed patch-wise cell-type proportions and assessed their correlation with Pearson correlation coefficient between those derived from the imputed transcriptomic data and CODEX data. This analysis demonstrates that STPAINTER preserves biologically plausible spatial organization of the TME.

Major lineages, including T cells, epithelial cells, and myeloid cells, were further subclustered and manually annotated via cluster-specific DEGs. The resulting cell type assignments were cross-validated with adjacent CODEX sections by calculating Pearson’s *r* of cell counts within matched 100µm×100µm regions.

### 4.6 Evaluation Metrics

To evaluate model performance, we employed a suite of metrics categorized into gene level and cluster level.

#### Gene Metrics

These metrics quantify the similarity between predicted gene expression 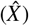 and ground-truth observations (*X*) across *n* cells.

- **Pearson correlation coefficient (PCC):** PCC measures the linear relationship between the predicted and observed expression profiles for each gene:

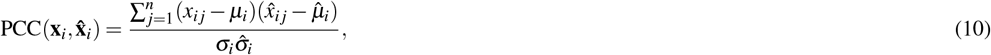

where *µ* and *σ* denote the mean and standard deviation of expression values, respectively.
- **Structural similarity index measure (SSIM):** SSIM assesses the preservation of spatial patterns by integrating luminance, contrast, and structural information:

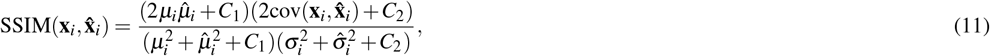

where cov(**x**_*i*_, 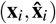) represents the covariance between **x**_*i*_ and 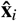, and *C*_1_,*C*_2_ are stabilization constants.
- **Root mean square error (RMSE):** RMSE quantifies the average prediction error magnitude. To account for cross-platform technical variation, it is calculated on *z*-score normalized expression values:

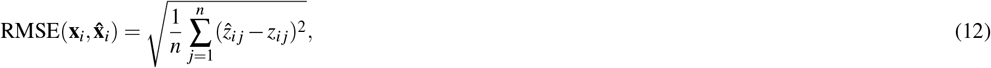

where *z*_*i j*_ and 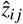 are normalized intensities. A lower RMSE indicates superior imputation accuracy.
- **Jaccard similarity (JS):** JS evaluates the similarity of spatial distributions, computed here as the Jensen–Shannon divergence between normalized spatial probability distributions **P**_*i*_ and 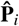:

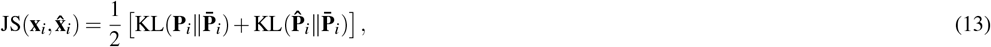

where 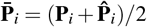 and KL denotes the Kullback–Leibler divergence.

#### Cluster Metrics

These metrics assess the model’s capacity to preserve the spatial organization of cell types and tissue domains.

- **Adjusted Rand Index (ARI):** ARI evaluates the similarity between the clustering of imputed data and ground-truth annotations, corrected for chance:

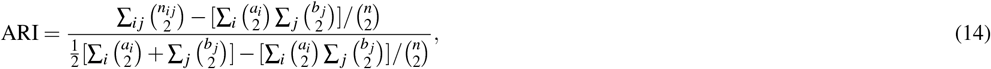

where *n*_*i j*_ is the number of shared elements between clusters, and *a*_*i*_, *b* _*j*_ are the marginal sums of the contingency table.
- **Adjusted Mutual Information (AMI):** AMI quantifies the information shared between two partitions, adjusted for the expected mutual information due to random chance:

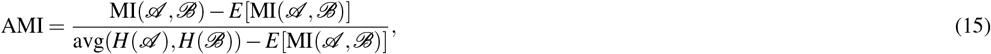

where MI is the mutual information, *H* is the entropy, and *E* represents the expectation.
- **Normalized Mutual Information (NMI):** NMI scales the mutual information score to a [0, 1] range based on the entropy of the partitions:

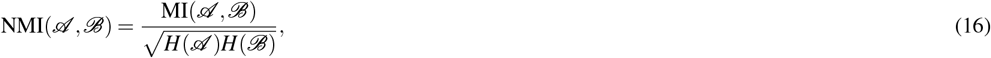

**Homogeneity (Homo):** Homogeneity measures the degree to which each cluster contains only members of a single class:

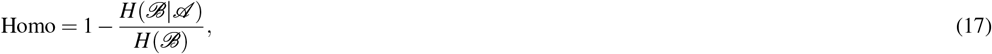

where *H*(*ℬ* | *𝒜*) is the conditional entropy of ground-truth categories *ℬ* given predicted assignments *𝒜*. A value of 1.0 indicates perfect homogeneity.

## Supporting information

Supplementary Information for stPainter

## Author contributions statement

Conceptualization: Y.H.Y. and Y.M.L.; Methodology: Y.H.Y. and Y.M.L.; Formal Analysis: Y.H.Y. and Y.M.L.; Software:Y.H.Y. and B.M.; Investigation: Y.H.Y. and Y.M.L.; Data Curation: Y.M.L.; Validation: Y.H.Y. and Y.M.L.; Visualization:Y.H.Y. and Y.M.L.; Writing – Original Draft: Y.H.Y. and Y.M.L.; Writing – Review & Editing: Y.H.Y., Y.M.L., K.Z., Z.X.Z., H.X.P., C.L.C., Q.L., and Y.C.; Supervision: K.Z.; Project Administration: K.Z.; Resources: K.Z., Y.C., L.S., and E.H.C.; Funding Acquisition: K.Z., Y.C., L.S., and E.H.C. All authors reviewed the manuscript.

## Data availability

The pan-cancer scRNA-seq datasets used for pretraining STPAINTER were obtained from publicly available studies deposited in the GEO dataset. Detailed accession numbers are provided in Table S1. Spatial transcriptomics datasets analyzed in this study include the SPATCH dataset (COAD, OV, and LIHC) as well as additional publicly available Xenium datasets (PRAD, NSCLC, and CESC). The SPATCH dataset is publicly available at https://spatch.pku-genomics.org^27^. Xenium 5k spatial transcriptomics datasets were obtained from the 10x Genomics public data repository, including datasets available at: https://www.10xgenomics.com/datasets/xenium-prime-ffpe-human-prostate, https://www.10xgenomics.com/datasets/xenium-human-lung-cancer-post-xenium-technote, and https://www.10xgenomics.com/datasets/xenium-prime-ffpe-human-cervical-cancer.

All processed data generated during this study are available from the corresponding author on a reasonable request.

## Code availability

All codes of proposed STPAINTER are published in https://github.com/yyh030806/stPainter.

## References

1. Gulati, G. S., D’Silva, J. P., Liu, Y., Wang, L. & Newman, A. M. Profiling cell identity and tissue architecture with single-cell and spatial transcriptomics. Nat. Rev. Mol. Cell Biol. 26, 11–31 (2025).

2. Liu, Y., Dai, Y. & Wang, L. Spatial omics at the forefront: emerging technologies, analytical innovations, and clinical applications. Cancer Cell (2025).

3. He, S. et al. High-plex imaging of rna and proteins at subcellular resolution in fixed tissue by spatial molecular imaging. Nat. biotechnology 40, 1794–1806 (2022).

4. Janesick, A. et al. High resolution mapping of the tumor microenvironment using integrated single-cell, spatial and in situ analysis. Nat. communications 14, 8353 (2023).

5. Tian, L., Chen, F. & Macosko, E. Z. Moving genomics into tissues. Nat. biotechnology 41, 773 (2022).

6. Ståhl, P. L. et al. Visualization and analysis of gene expression in tissue sections by spatial transcriptomics. Science 353, 78–82 (2016).

7. Asp, M. et al. A spatiotemporal organ-wide gene expression and cell atlas of the developing human heart. Cell 179, 1647–1660 (2019).

8. Moncada, R. et al. Integrating microarray-based spatial transcriptomics and single-cell rna-seq reveals tissue architecture in pancreatic ductal adenocarcinomas. Nat. biotechnology 38, 333–342 (2020).

9. Kwok, A. W. C. et al. Denoising image-based spatial transcriptomics data with denoist. bioRxiv 2025–11 (2025).

10. Bilous, M. et al. From transcripts to cells: Dissecting sensitivity, signal contamination, and specificity in xenium spatial transcriptomics. bioRxiv 2025–04 (2025).

11. Hao, Y. et al. Dictionary learning for integrative, multimodal and scalable single-cell analysis. Nat. biotechnology 42, 293–304 (2024).

12. Biancalani, T. et al. Deep learning and alignment of spatially resolved single-cell transcriptomes with tangram. Nat. methods 18, 1352–1362 (2021).

13. Abdelaal, T., Mourragui, S., Mahfouz, A. & Reinders, M. J. Spage: spatial gene enhancement using scrna-seq. Nucleic acids research 48, e107–e107 (2020).

14. Shengquan, C., Boheng, Z., Xiaoyang, C., Xuegong, Z. & Rui, J. stplus: a reference-based method for the accurate enhancement of spatial transcriptomics. Bioinformatics 37, i299–i307 (2021).

15. Moriel, N. et al. Novosparc: flexible spatial reconstruction of single-cell gene expression with optimal transport. Nat. protocols 16, 4177–4200 (2021).

16. Cang, Z. & Nie, Q. Inferring spatial and signaling relationships between cells from single cell transcriptomic data. Nat. communications 11, 2084 (2020).

17. Welch, J. D. et al. Single-cell multi-omic integration compares and contrasts features of brain cell identity. Cell 177, 1873–1887 (2019).

18. Lopez, R. et al. A joint model of unpaired data from scrna-seq and spatial transcriptomics for imputing missing gene expression measurements. arXiv preprint arXiv:1905.02269 (2019).

19. Wan, X. et al. Integrating spatial and single-cell transcriptomics data using deep generative models with spatialscope. Nat. Commun. 14, 7848 (2023).

20. Sun, E. D., Ma, R. & Zou, J. Sprite: improving spatial gene expression imputation with gene and cell networks. Bioinformatics 40, i521–i528 (2024).

21. Li, K., Li, J., Tao, Y. & Wang, F. stdiff: a diffusion model for imputing spatial transcriptomics through single-cell transcriptomics. Briefings Bioinforma. 25 (2024).

22. Li, B. et al. Spaim: Single-cell spatial transcriptomics imputation via style transfer. bioRxiv (2025).

23. Rombach, R., Blattmann, A., Lorenz, D., Esser, P. & Ommer, B. High-resolution image synthesis with latent diffusion models. In Proceedings of the IEEE/CVF conference on computer vision and pattern recognition, 10684–10695 (2022).

24. Song, Y. et al. Score-based generative modeling through stochastic differential equations. arXiv preprint arXiv:2011.13456 (2020).

25. Marco Salas, S. et al. Optimizing xenium in situ data utility by quality assessment and best-practice analysis workflows. Nat. Methods 1–11 (2025).

26. Chang, J. et al. Single-cell multi-stage spatial evolutional map of esophageal carcinogenesis. Cancer Cell 43, 380–397 (2025).

27. Ren, P. et al. Systematic benchmarking of high-throughput subcellular spatial transcriptomics platforms across human tumors. Nat. Commun. 16, 9232 (2025).

28. Zheng, L. et al. Pan-cancer single-cell landscape of tumor-infiltrating t cells. Science 374, abe6474 (2021).

29. You, S. et al. Lymphatic-localized treg-mregdc crosstalk limits antigen trafficking and restrains anti-tumor immunity. Cancer cell 42, 1415–1433 (2024).

30. Loi, P. et al. Epigenetic and oncogenic inhibitors cooperatively drive differentiation and kill kras-mutant colorectal cancers. Cancer Discov. 14, 2430–2449 (2024).

31. Brooks, D. L. P. et al. Itga6 is directly regulated by hypoxia-inducible factors and enriches for cancer stem cell activity and invasion in metastatic breast cancer models. Mol. cancer 15, 26 (2016).

32. Van der Flier, L. G., Haegebarth, A., Stange, D. E., Van de Wetering, M. & Clevers, H. Olfm4 is a robust marker for stem cells in human intestine and marks a subset of colorectal cancer cells. Gastroenterology 137, 15–17 (2009).

33. Marteau, V. et al. Single-cell integration and multi-modal profiling reveals phenotypes and spatial organization of neutrophils in colorectal cancer. Cancer Cell 44, 146–165 (2026).

34. Lopez, R., Regier, J., Cole, M. B., Jordan, M. I. & Yosef, N. Deep generative modeling for single-cell transcriptomics. Nat. methods 15, 1053–1058 (2018).

35. Denisenko, E. et al. Systematic assessment of tissue dissociation and storage biases in single-cell and single-nucleus rna-seq workflows. Genome biology 21, 130 (2020).

36. van den Brink, S. C. et al. Single-cell sequencing reveals dissociation-induced gene expression in tissue subpopulations. Nat. methods 14, 935–936 (2017).

37. Peebles, W. & Xie, S. Scalable diffusion models with transformers. In Proceedings of the IEEE/CVF international conference on computer vision, 4195–4205 (2023).

38. Anderson, B. D. Reverse-time diffusion equation models. Stoch. Process. their Appl. 12, 313–326 (1982).

39. Wolf, F. A., Angerer, P. & Theis, F. J. Scanpy: large-scale single-cell gene expression data analysis. Genome biology 19, 15 (2018).

