## Supplementary Information for stPainter for "Enhancing Pan-cancer Spatial Transcriptomics at Single-cell Resolution with STPAINTER"

#### S1 Overview of the Pan-cancer scRNA-seq Atlas

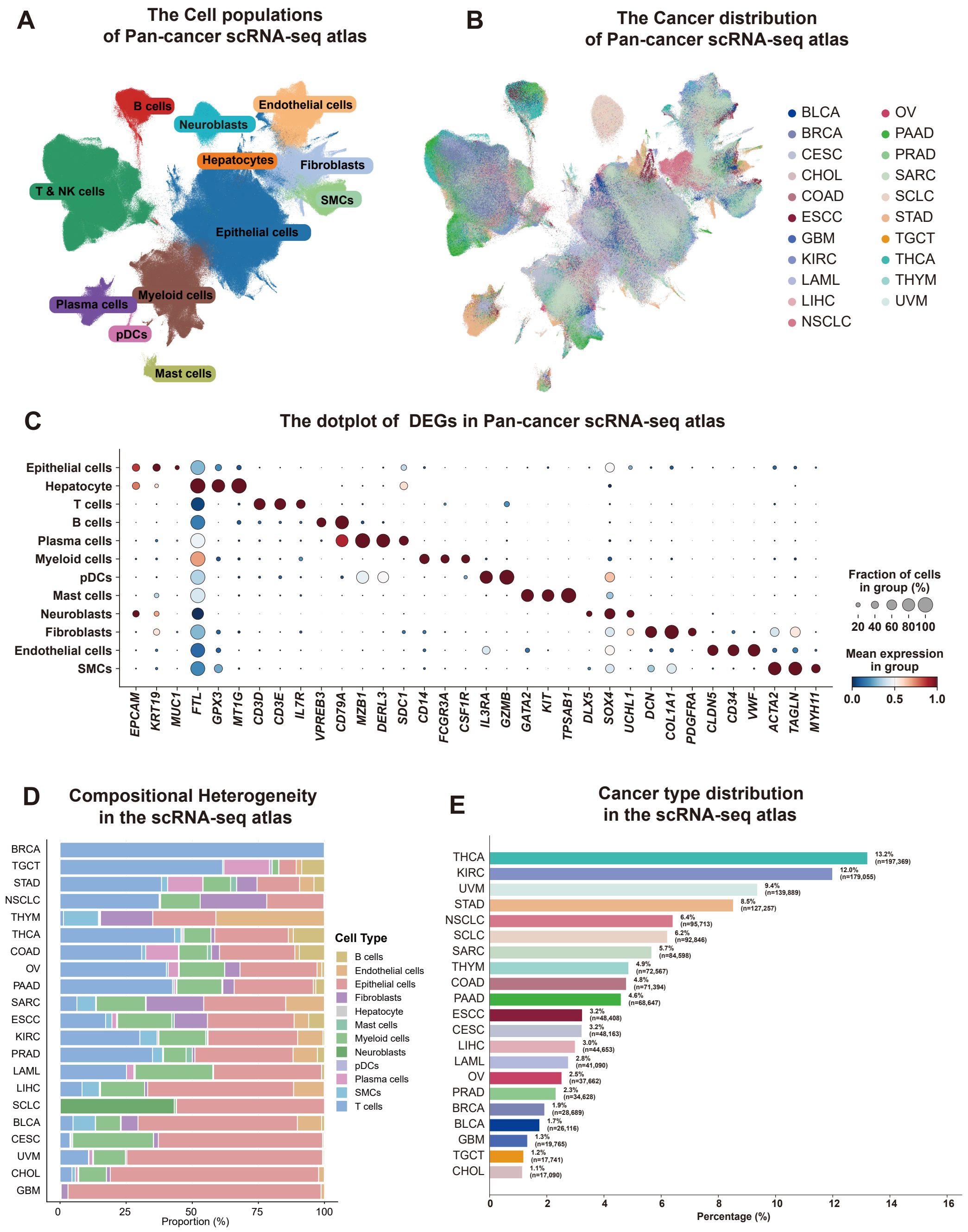


**Fig. S1. Details of the pan-cancer scRNA-seq atlas. a**, UMAP of all major cell populations of Pan-cancer scRNA-seq atlas. **b**, UMAP of all cancer types distributions of Pan-cancer scRNA-seq atlas. **c**, dotplot of DEGs in Pan-cancer scRNA-seq atlas. **d**, the compositional heterogeneity in Pan-cancer scRNA-seq atlas. **e**, the percentage of cell populations in different cancer types of Pan-cancer scRNA-seq atlas.

To construct a comprehensive reference for the STPAINTER framework, we aggregated a pan-cancer single-cell RNA sequencing (scRNA-seq) atlas from publicly available studies in the Gene Expression Omnibus (GEO) database. As detailed in **Table**

**S1**, this dataset encompasses 21 diverse cancer types. We employed scVI for batch-corrected integration, which successfully resolved distinct cell populations (Fig. S1a) validated by specific canonical marker expression patterns (Fig. S1c). These findings demonstrate the high fidelity of the single-cell atlas, establishing it as a robust reference for pretraining. Furthermore, the datasets exhibited diverse cellular distributions and cancer-type-specific tumor microenvironment heterogeneity (Fig. S1b, d), providing a comprehensive and diverse foundation for pan-cancer model training.

| Abbr. | Cancer Type | Source | Journal | Platform |
|---|---|---|---|---|
| LAML | Acute Myeloid Leukemia | GSE116256 | Cell | 10X Genomics |
| BLCA | Bladder Urothelial Carcinoma | GSE222315 | Cancer Cell | 10X Genomics |
| BRCA | Breast invasive carcinoma | GSE114724 | Cell | 10X Genomics |
| CESC | Cervical squamous cell carcinoma and endocervical adenocarcinoma | GSE144240 | Cell | 10X Genomics |
| CHOL | Cholangiocarcinoma | GSE138709 | Journal of Hepatology | 10X Genomics |
| COAD | Colon adenocarcinoma | GSE166555 | EMBO Mol Med | 10X Genomics |
| COAD | Colon adenocarcinoma | GSE188711 | JCI Insight | 10X Genomics |
| ESCC | esophageal squamous cell carcinoma | GSE188900 | Signal Transduct Target Ther | 10X Genomics |
| GBM | Glioblastoma multiforme | GSE193884 | Cancer Cell | 10X Genomics |
| KIRC | Kidney renal clear cell carcinoma | GSE207493 | Cancer Res | 10X Genomics |
| LIHC | Liver hepatocellular carcinoma | GSE112271 | Nat Commun | 10X Genomics |
| NSCLC | non-small cell lung cancer | GSE162498 | Sci Immunol | 10X Genomics |
| NSCLC | non-small cell lung cancer | GSE233203 | MedComm | 10X Genomics |
| SCLC | small cell lung cancer | GSE138474 | Cancer Cell | 10X Genomics |
| OV | Ovarian serous cystadenocarcinoma | GSE184880 | Clin Cancer Res | 10X Genomics |
| PAAD | Pancreatic adenocarcinoma | GSE154778 | Genome Med | 10X Genomics |
| PAAD | Pancreatic adenocarcinoma | GSE263733 | Mol Cancer | 10X Genomics |
| PRAD | Prostate adenocarcinoma | GSE193337 | Mol Cancer | 10X Genomics |
| SARC | Sarcoma | GSE212527 | Nat Cancer | 10X Genomics |
| STAD | Stomach adenocarcinoma | GSE183904 | Cancer Discov | 10X Genomics |
| TGCT | Testicular Germ Cell Tumors | GSE228501 | Nat Commun | 10X Genomics |
| THYM | Thymoma | GSE212527 | Nat Commun | 10X Genomics |
| THCA | Thyroid carcinoma | GSE184362 | Nat Commun | 10X Genomics |
| UVM | Uveal Melanoma | GSE139829 | Nat Commun | 10X Genomics |

**Table S1.** Metadata of Pan-cancer dataset from GEO database

### S2 Comparative Analysis of Existing Computational Enhancement Methods

#### S2.1 Alignment-based Methods

Alignment-based methods reconstruct spatial context by establishing statistical or geometric correspondence between dissociated scRNA-seq profiles and a reference spatial scaffold. These approaches typically utilize feature matching, where a subset of marker genes shared across both modalities guides the probabilistic mapping of single cells onto discrete spatial locations. By optimizing an alignment cost function formulated through optimal transport or manifold alignment, these tools project high-resolution transcriptomic information onto histological structures. This paradigm is particularly effective for genome-wide imputation and deconvolving coarse-resolution spatial data into single-cell resolution.

Tangram[12] is a deep-learning framework that maps sc/snRNA-seq data onto spatial supports by optimizing a probabilistic alignment matrix. The algorithm treats single-cell profiles as essential components and rearranges them to maximize the spatial correlation of shared genes between modalities. Its objective function primarily utilizes Kullback-Leibler divergence to match cell-density distributions and cosine similarity to align gene expression patterns. Beyond simple mapping, Tangram can project the entire transcriptome onto spatial coordinates, effectively expanding targeted gene panels to genome-wide resolution, correcting low-quality spatial measurements, and deconvolving low-resolution data to single-cell resolution.

SpaOTsc[16] is a computational method that utilizes structured and unbalanced optimal transport to recover spatial properties for scRNA-seq data. By mapping single cells to a reference spatial measurement of a limited gene set, the algorithm establishes a spatial metric at single-cell resolution. This metric enables the reconstruction of spatial gene expression patterns, the identification of spatially localized cell subclusters, and the inference of space-constrained cell-cell communications mediated by ligand-receptor interactions. Furthermore, SpaOTsc employs partial information decomposition to quantify intercellular gene-gene regulatory information flows, allowing for the estimation of signaling spatial ranges and potential cross-cell genetic regulations.

NovoSpaRc[15] is a computational framework that probabilistically assigns single cells to tissue locations using an optimal transport framework. The method is built upon the structural correspondence hypothesis, which posits that cells in physical proximity share similar gene expression profiles. By minimizing the deviation between expression-based distances and physical target space coordinates, NovoSpaRc can reconstruct tissue organization *de novo*. While the model can operate solely on scRNA-seq data, its performance is significantly enhanced by incorporating a reference atlas of marker genes when available.

##### S2.2 Generative-based Methods

Generative-based methods leverage deep generative architectures to explicitly model the underlying probability distribution of cellular gene expression. Unlike rigid alignment strategies, these frameworks internalize complex biological patterns to synthesize or reconstruct spatial profiles. By accounting for modality-specific noise and technical biases through latent variable modeling, generative approaches provide a robust foundation for denoising, imputation, and identifying continuous biological gradients across diverse tissue architectures.

gimVI[18] is a generative framework designed to integrate scRNA-seq and ST data by leveraging variational autoencoders. The model learns a joint latent representation of cells from both modalities, enabling the imputation of genes not originally measured in the spatial dataset. By modeling the data using a deep generative architecture, gimVI accounts for technical noise and modality-specific biases while preserving biological signals. The framework allows for the probabilistic transfer of information across datasets, providing a robust approach for reconstructing high-resolution spatial gene expression profiles and identifying spatially organized cellular states.

stDiff[21] is a generative framework that utilizes a conditional diffusion model to enhance the quality and resolution of ST data. By incorporating scRNA-seq as a reference, stDiff learns to reconstruct the spatial distribution of the whole transcriptome through a reverse diffusion process. The model treats spatial imputation as a conditional generation task, where the diffusion architecture effectively captures intricate spatial gradients and recovers missing biological signals in sparse datasets. This approach is particularly adept at preserving localized expression patterns and correcting technical dropouts, facilitating high-fidelity spatial analysis at single-cell resolution.

SpaIM[22] is a style transfer learning model designed to predict unmeasured gene expressions in ST profiles by leveraging scRNA-seq references. The core architecture focuses on disentangling shared biological content from modality-specific styles, effectively integrating the rich transcriptomic coverage of scRNA-seq with the spatial context of ST data. By employing a style transfer paradigm, SpaIM accounts for the technical discrepancies between sequencing- and imaging-based platforms, enabling robust cross-modality information transfer. This framework enhances downstream applications, including ligand-receptor interaction inference and spatial domain characterization.

#### S3 Technical Implementation and Hyperparameter Configurations

##### S3.1 Architecture Hyperparameters of the VAE

The VAE within the STPAINTER framework serves as the foundational component for dimensionality reduction and noise filtering[34]. Its primary objective is to map high-dimensional, sparse gene expression profiles into a compact and structured latent manifold, $\mathbf{z}$, while disentangling biological signals from technical nuisance factors.

The encoder network, $\text{Encoder}_{\mathbf{z}}(\mathbf{x}, \mathbf{c})$, processes the input gene expression vector $\mathbf{x}$ and a one-hot encoded cancer-type vector $\mathbf{c}$ to estimate the parameters of the latent distribution. To account for technical variation in sequencing depth, the model additionally predicts a scalar library size factor $l \sim \text{LogNormal}(\mu_l, \sigma_l^2)$. The decoder network, $\text{Decoder}(\mathbf{z}, \mathbf{c})$, then maps the sampled latent variables back to the gene space, outputting the parameters $(\rho, \pi)$ of a ZINB distribution to model the inherent sparsity and over-dispersion of the data.

We implemented two model variants, STPAINTER-50 and STPAINTER-100, which differ solely in the dimension of their latent space to accommodate varying levels of biological complexity. The detailed architectural parameters are summarized in Table. S2.

##### S3.2 Architecture Hyperparameters of the GiT

The GiT serves as the core generative module of STPAINTER, designed to model the complex probability distribution of cellular states within the learned latent manifold[23]. The architecture is based on the Scalable Diffusion Transformer framework[37], specifically optimized for processing 1D latent transcriptomic embeddings.

| Parameter | Description | stPainter-50 | stPainter-100 |
|---|---|---|---|
| Input Size ($G$) | Number of target genes in the pan-cancer atlas | 10,000 | 10,000 |
| Latent Size ($L$) | Dimension of the biological latent manifold $z$ | 50 | 100 |
| Hidden Size | Number of neurons in each hidden layer | 256 | 256 |
| Number of Layers | Depth of the encoder and decoder networks | 4 | 4 |
| Conditional Classes ($K$) | Number of cancer types used for conditioning | 21 | 21 |
| Output Model | Probabilistic likelihood distribution for reconstruction | ZINB | ZINB |
| Activation Function | Non-linear activation used between hidden layers | LeakyReLU | LeakyReLU |

**Table S2.** Architectural Specifications for the VAE Components

The input latent vector $\mathbf{z}(t)$ is treated as a sequence of $L$ tokens, where $L$ corresponds to the dimension of the VAE latent space ($L = 50$ or 100). Each token is projected into a hidden dimension $D$ and augmented with 1D learnable positional embeddings to preserve topological semantics. The model employs the adaLN-Zero mechanism to rigorously condition the denoising process on the cancer-type context $\mathbf{c}$ and the diffusion timestep $t$. The specific configurations for the GiT are detailed in Table. S3.

| Parameter | Description | Value |
|---|---|---|
| Depth | Number of stacked Transformer blocks | 8 |
| Hidden Size ($D$) | Dimension of the transformer hidden states | 128 |
| Number of Heads | Heads in MSA | 8 |
| MLP Ratio | Expansion factor in the FFN | 4.0 |
| Patch Size | Size of the input patch (feature-wise tokenization) | 1 |
| Conditioning | Modulation method for external priors | adaLN-Zero |
| Positional Embedding | Type of spatial/topological encoding | Learnable 1D |

**Table S3.** Architectural Specifications for the GiT Components

##### S3.3 Training Strategy and Hyperparameters

The training of stPainter is implemented as a two-stage sequential optimization process to ensure the stability of the generative manifold[23]. In the first stage, the VAE is trained on the pan-cancer scRNA-seq atlas to maximize the ELBO, balancing ZINB reconstruction fidelity with Kullback-Leibler (KL) divergence regularization[34]. In the second stage, with the VAE weights frozen, the GiT is trained to model the conditional distribution of the latent cellular states[24].

Both modules were trained on the integrated pan-cancer dataset using the AdamW optimizer with a learning rate of $1 \times 10^{-4}$. A class dropout probability of 0.1 was applied during GiT training to support robust conditional generation. The training epochs for the VAE and GiT were set to 200 and 1,000, respectively, ensuring thorough learning of the universal cellular prior. The specific hyperparameter configurations used during the pretraining phase are summarized in Table. S4.

| Parameter | Description | VAE | GiT |
|---|---|---|---|
| Batch Size | Number of samples per training step | 1,024 | 1,024 |
| Learning Rate | Initial step size for the optimizer | $1 \times 10^{-4}$ | $1 \times 10^{-4}$ |
| Training Epochs | Total number of passes over the dataset | 200 | 1,000 |
| Optimizer | Algorithm used for weight updates | AdamW | AdamW |
| Loss Function | Objective used for optimization | ELBO | MSE |
| Class Dropout | Probability for classifier-free guidance | N/A | 0.1 |

**Table S4.** Training Hyperparameters for VAE and GiT Modules

##### S3.4 Sampling and Inference Hyperparameters

The inference phase of STPAINTER utilizes a guided generative process to transform partial spatial measurements into high-fidelity, genome-wide profiles[?]. This is achieved by initializing the reverse diffusion process from a controlled corrupted state rather than pure noise. The balance between preserving the original spatial observations and leveraging the learned pan-cancer generative prior is primarily governed by the forward perturbation level ($t_0$).

During inference, we employ a SDE solver[24] to iteratively refine the latent embeddings. To ensure the generated profiles conform to the desired biological context, we incorporate Classifier-Free Guidance (CFG) during the denoising steps. The specific hyperparameter configurations for the sampling and transport processes, including the SDE integration method and the velocity prediction target, are detailed in Table. S5.

| Parameter | Description | Value |
|---|---|---|
| Sampling Mode | Numerical framework for reverse diffusion | SDE |
| Forward Level ($t_0$) | Initial noise level added to ST measurements | 0.9 |
| Sampling Steps | Number of iterative refinement steps | 30 |
| CFG Scale | Strength of Classifier-Free Guidance | 1.0 |
| SDE Solver | Numerical method for stochastic integration | Euler |
| Path Type | Mathematical form of the probability path | Linear |
| Prediction Type | Target of the network output | Velocity |
| Last Step Strategy | Final correction method for gene reconstruction | Mean |

**Table S5.** Hyperparameters for Diffusion Sampling and Inference

#### S4 Performance Benchmarking and Generalizability Validation

##### S4.1 Quantitative Results in Diverse Tumor Tissues (OV and LIHC)

Quantitative benchmarking on the OV dataset across varying numbers of HVGs confirms the robustness of STPAINTER. On the OV dataset, both STPAINTER-100 and STPAINTER-50 consistently outperformed baseline methods across multiple evaluation metrics. As the number of imputed genes scaled from 10 to 300, STPAINTER-50 achieved a superior PCC. This metric reached 0.37 for the top 100 genes, surpassing the performance of leading baselines such as gimVI (PCC = 0.34) and Tangram (PCC = 0.32). The advantage in structural fidelity was similarly pronounced. STPAINTER-50 attained an SSIM of 0.27 at the 100 HVG scale, while gimVI and Tangram yielded significantly lower scores of 0.18 and 0.07, respectively. The error metrics further support the high-fidelity recovery of gene expression in this tissue type. At the 100 HVG scale, STPAINTER-100 achieved a low global error with an RMSE of 1.12 and a JS of 0.45. These results demonstrate that the imputed profiles accurately capture both the absolute abundance and the statistical distribution of the ground-truth transcripts. Similar to observations in the COAD dataset, the performance of STPAINTER remained stable as the complexity of the gene panel increased, highlighting the effectiveness of the universal cellular prior in regularizing the generation process. Unsupervised clustering analysis on the OV dataset further illustrates the capacity of STPAINTER to resolve distinct spatial domains. STPAINTER-50 achieved an ARI of 0.80, an AMI of 0.63, a Homogeneity score of 0.57, and an NMI of 0.63. While gimVI exhibited a high ARI of 0.86, STPAINTER maintained competitive performance across all metrics. Specifically, STPAINTER-100 yielded a Homogeneity score of 0.62, which was the highest among all tested methods, indicating superior biological specificity in cluster assignments. In contrast, other generative models such as SpaIM showed significantly lower performance, with an ARI of 0.17. These consistent gains across metrics suggest that STPAINTER reconstructs a high-fidelity cellular manifold that is essential for accurate tissue segmentation in complex cancer landscapes.

The evaluation on the LIHC dataset further underscores the generalizability and robustness of STPAINTER across diverse cancer histologies (Fig. S2c, d). Quantitative benchmarking of gene expression recovery demonstrates that STPAINTER variants consistently outperform all baseline methods across varying genomic scales (Fig. S2c). As the number of imputed genes increased from 10 to 300, STPAINTER-50 achieved a superior PCC, peaking at 0.35 for the top 100 genes, compared to the best generative baseline gimVI (PCC = 0.31) and alignment-based Tangram (PCC = 0.30). Structural fidelity was similarly enhanced, with STPAINTER-50 maintaining an SSIM of 0.39 at the 100 HVG scale, while alignment methods like Tangram and NovoSpaRc yielded scores below 0.12. Furthermore, STPAINTER variants minimized global error and statistical divergence, achieving the lowest RMSE (1.11) and JS (0.45) at the 100 HVG scale. This stable performance across increasing gene complexity highlights the effectiveness of the learned universal cellular prior in capturing high-fidelity expression manifolds. Beyond gene recovery, STPAINTER effectively resolved the spatial architecture of LIHC through high-resolution tissue segmentation (Fig. S2d). Quantitative analysis of clustering performance shows that STPAINTER-100 achieved the highest scores in structural

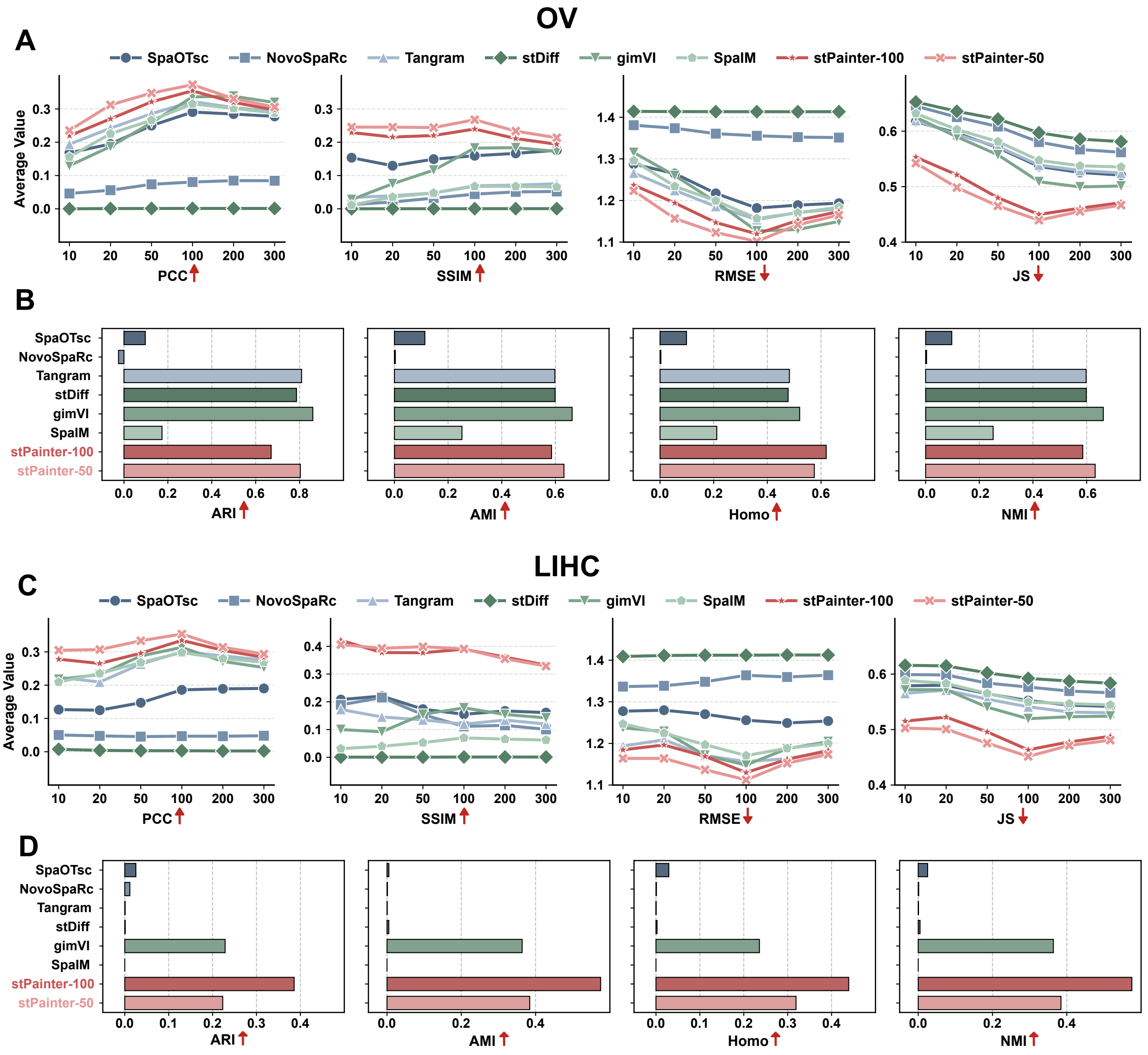


**Fig. S2. Detailed benchmarking of gene expression recovery and spatial domain identification on OV and LIHC datasets. a,** Quantitative assessment of gene expression recovery on the Ovarian Cancer (OV) dataset across varying genomic scales (10 to 300 highly variable genes, HVGs). Performance is measured by Pearson Correlation Coefficient (PCC), Structural Similarity Index Measure (SSIM), Root Mean Square Error (RMSE), and Jensen-Shannon divergence (JS). **b,** Benchmarking of unsupervised clustering performance on the OV dataset using four metrics: Adjusted Rand Index (ARI), Adjusted Mutual Information (AMI), Homogeneity (Homo), and Normalized Mutual Information (NMI). **c,** Quantitative evaluation of gene expression recovery on the Liver Hepatocellular Carcinoma (LIHC) dataset across different HVG scales. **d,** Evaluation of structural resolution and tissue segmentation accuracy on the LIHC dataset across clustering metrics. For both cancer types, STPAINTER variants consistently demonstrate superior fidelity in reconstructing transcriptomic manifolds compared to baseline alignment-based and generative methods.

resolution, with an ARI of 0.39, AMI of 0.58, and NMI of 0.58. Notably, STPAINTER-100 reached a Homogeneity score of 0.44, significantly exceeding baseline generative models such as gimVI (Homo = 0.24) and SpaIM (Homo = 0). In contrast, traditional alignment-based tools like Tangram (ARI = 0.00) and NovoSpaRc (ARI = 0.01) failed to identify biologically meaningful spatial domains in the LIHC landscape. These results demonstrate that the latent representations generated by

STPAINTER successfully preserve fine-grained cellular heterogeneity, facilitating accurate spatial domain identification even in complex tumor microenvironments.

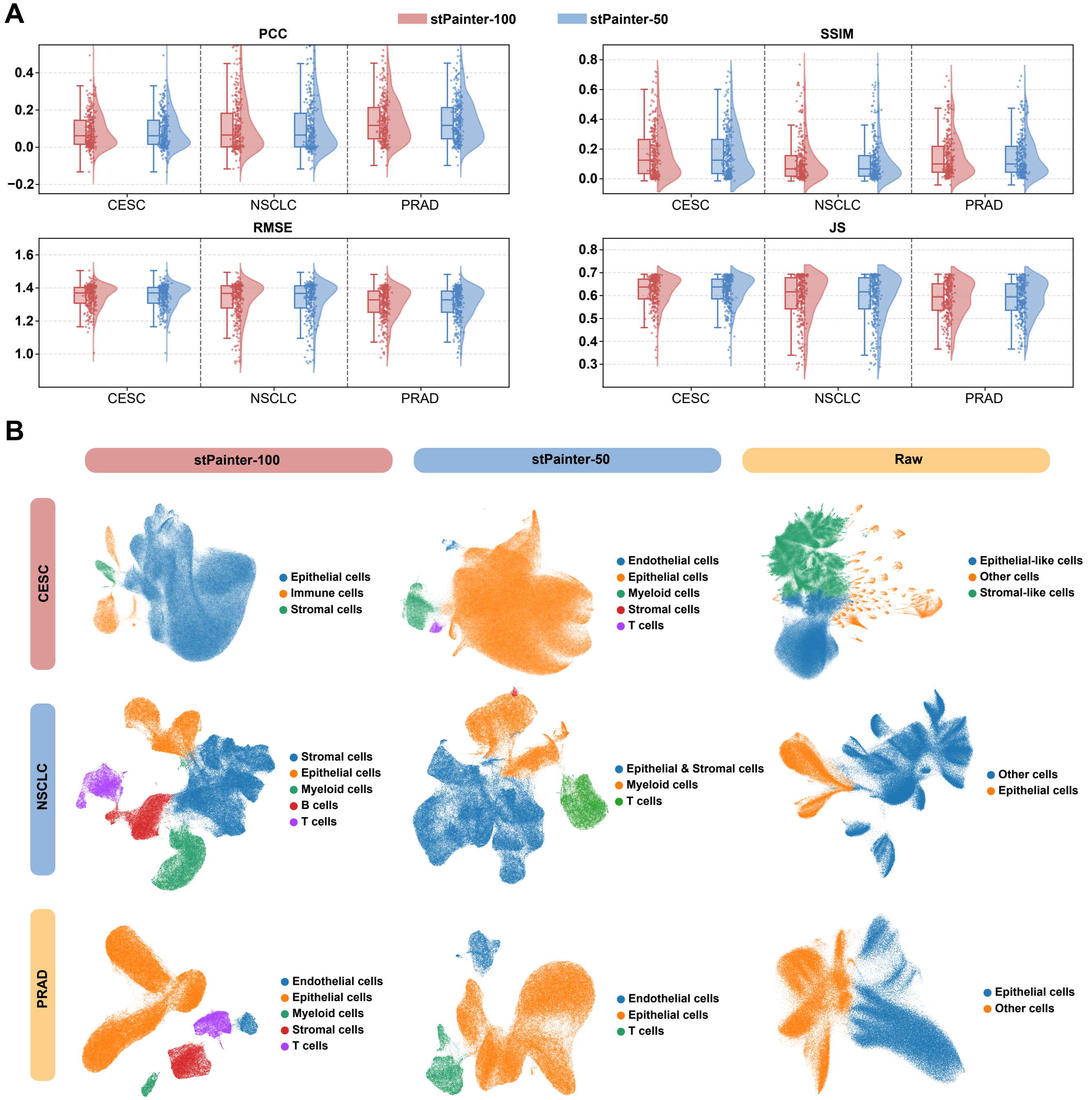


**Fig. S3. Zero-shot generalization and topological preservation on datasets without matched scRNA-seq references. a,** Performance benchmarking of STPAINTER on CESC, NSCLC, and PRAD datasets where tissue-specific single-cell references were unavailable. Gene-level recovery metrics (PCC, SSIM, RMSE, and JS) demonstrate the model's ability to leverage the universal cellular prior for high-fidelity imputation across unseen histological landscapes. **b,** UMAP visualizations comparing the topological structures of cell populations derived from raw spatial measurements, STPAINTER latent representations, and baseline methods. STPAINTER effectively resolves distinct cellular clusters and biological gradients that are obscured by sparsity and technical noise in the raw data or artifacts in competing generative workflows.

##### S4.2 Zero-shot Enhancement on Unseen Pathological Landscapes

To further evaluate the robustness and universal applicability of stPainter, we extended our benchmarking to three additional spatial transcriptomics datasets, namely Cervical Squamous Cell Carcinoma (CESC), Non-Small Cell Lung Cancer (NSCLC), and Prostate Adenocarcinoma (PRAD), for which matched scRNA-seq reference data were unavailable. This scenario rigorously tests the model's zero-shot generalization capabilities, as it must rely solely on the universal cellular manifolds learned from the massive pan-cancer atlas during the pretraining phase.

As illustrated in Fig. S3a, both stPainter-100 and stPainter-50 variants demonstrated consistent high-fidelity recovery across varying genomic scales ranging from 10 to 300 highly variable genes. Quantitatively, on the PRAD dataset, stPainter-100 achieved a peak PCC of 0.125 and a SSIM of 0.268. Similar performance stability was observed in the CESC dataset, where the PCC reached 0.086 and the SSIM remained above 0.279 for sparse gene panels. In the NSCLC landscape, despite the inherent technical noise, the model maintained low global errors with an RMSE of 1.371 and stable statistical divergence with a JS of 0.630. These findings substantiate that the learned universal cellular prior effectively regularizes the imputation process, enabling high-fidelity spatial gene expression recovery even in the absence of dataset-specific training or matched single-cell references.

Given the inherent noise and limited resolution of the raw Xenium data in these extended datasets, PCA-based clustering proved unreliable for establishing a ground truth or robust baseline. Consequently, we omitted quantitative clustering metrics for this analysis and instead relied on qualitative assessment at a consistent coarse resolution (Fig. S3b).

In the NSCLC and PRAD datasets, stPainter-100 demonstrated superior discriminative power compared to stPainter-50. Specifically, the higher-dimensional latent space allowed for the fine-grained delineation of immune subpopulations in PRAD and the precise distinction of stromal subsets in NSCLC. Conversely, the CESC dataset exhibited the poorest quality in its raw state; with the exception of epithelial cells, the raw data yielded scattered, biologically incoherent patterns on the UMAP embedding, indicative of severe signal sparsity. In this scenario, the lower-dimensional stPainter-50 outperformed stPainter-100, likely because the expanded latent space of the latter captured excessive technical noise, thereby compromising cluster separation. Nevertheless, both stPainter variants successfully reconstructed defined cellular clusters with significantly improved biological interpretability compared to the raw data.

##### S4.3 Ablation Study of Settings

To evaluate the sensitivity of stPainter to different sampling and guidance configurations, we conducted systematic ablation experiments focusing on the number of iterative refinement steps and the conditioning strength of the generative process across various genomic scales. As illustrated in Fig. S4, these parameters collectively determine the balance between computational efficiency and the biological fidelity of the imputed profiles.

Analysis of the performance across genomic scales in Fig. S4a reveals that for stPainter-100, the gene-level fidelity remains remarkably stable as iterative steps increase. For instance, at the 100 highly variable genes HVGs scale, stPainter-100 achieved a PCC of 0.316 and a SSIM of 0.302 with 10 steps, which maintained consistent levels through to 50 steps. However, the resolution of spatial domains shows a stronger dependency on the refinement duration. As shown in Fig. S4b, for the stPainter-100 variant, extending the process from 10 to 30 steps significantly improved the ARI from 0.786 to 0.847 and the NMI from 0.638 to 0.696. The clustering performance plateaued beyond 30 steps, with an ARI of 0.848 at 50 iterations, suggesting that 30 steps provide an optimal threshold for capturing fine-grained tissue architectures[24].

The biological specificity of the generation is further modulated by the intensity of the cancer-type guidance across the target gene panels. The quantitative metrics in Fig. S4c demonstrate that increasing guidance intensity generally leads to a decline in gene-level recovery accuracy. For stPainter-100 at 100 HVGs, the PCC decreased from 0.315 at 1.0 intensity to 0.260 at 2.5 intensity. Quantitative analysis of the spatial resolution in Fig. S4d reveals distinct behaviors between model variants. For stPainter-100, the peak structural resolution was observed at a guidance intensity of 1.0, yielding an ARI of 0.847 and a Homo score of 0.630. Higher conditioning levels resulted in a sharp reduction in clustering accuracy for this variant, with the ARI falling to 0.459 at 2.5 intensity. In contrast, the stPainter-50 variant exhibited a peak ARI of 0.810 at a guidance intensity of 2.0 before declining[37]. These findings indicate that while appropriate conditioning aligns the imputed data with specific malignant contexts, excessive intensity can introduce generative artifacts that obscure underlying biological signals.

##### S4.4 Computational Scalability and Inference Dynamics

To assess the practical deployability of stPainter, we conducted a comprehensive benchmarking of computational resources across three cancer datasets (COAD, LIHC, and OV), specifically comparing against GPU-accelerated baselines. As illustrated in Fig. S5a, we evaluated the running time and peak memory usage of stPainter variants (stPainter-100 and stPainter-50) relative to the alignment-based method Tangram and generative models including stDiff, gimVI, and SpaIM. Despite the

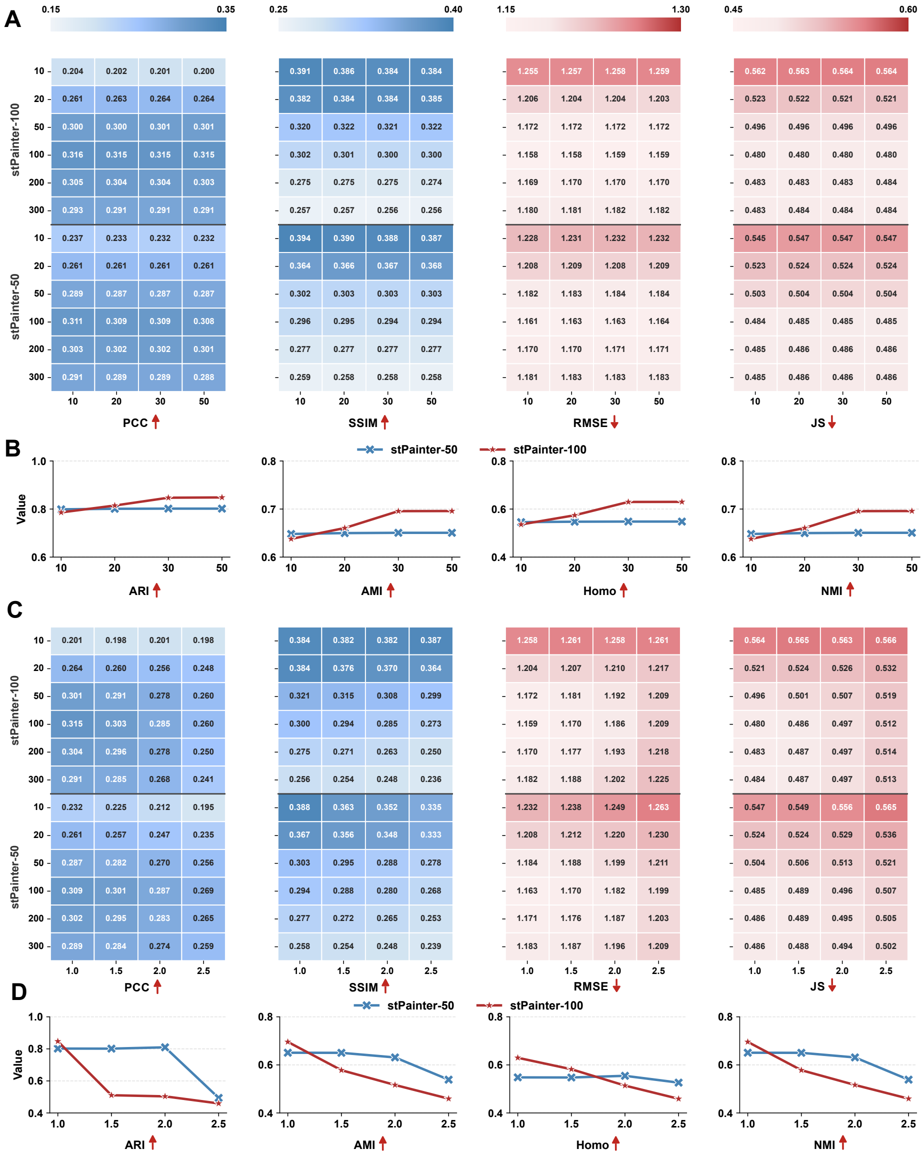


**Fig. S4. Ablation analysis of sampling dynamics and guidance intensity across genomic scales. a-b,** Impact of iterative refinement steps (10, 20, 30, and 50 iterations) on model performance. **a,** Quantitative metrics of gene-level fidelity (PCC, SSIM, RMSE, and JS) across varying HVG scales (10 to 300), showing the stability of transcriptomic recovery as a function of sampling duration. **b,** Influence of refinement steps on clustering resolution (ARI, AMI, Homo, and NMI), illustrating that 30 iterations represent an optimal threshold for capturing fine-grained tissue architectures. **c-d,** Sensitivity of STPAINTER variants to the intensity of cancer-type guidance (scales 1.0, 1.5, 2.0, and 2.5). **c,** Gene-level recovery accuracy across different guidance levels, where excessive conditioning may lead to a decline in global metrics. **d,** Cluster-level resolution performance, demonstrating that while STPAINTER-100 achieves peak structural resolution at lower guidance intensities, STPAINTER-50 exhibits higher tolerance to conditioning for optimal domain identification.

inherent computational intensity of iterative diffusion processes, STPAINTER demonstrates remarkable efficiency, performing comparably to or outperforming these GPU-based benchmarks. This scalability stems from our distinct two-stage architecture: by shifting the computationally expensive diffusion modeling from the high-dimensional gene space to a compressed latent manifold ($L = 50$ or 100), STPAINTER significantly reduces the GPU memory overhead and inference latency. Notably, it achieves a substantial speedup compared to pixel-space diffusion methods like stDiff, while maintaining a resource footprint competitive with other generative baselines. The logarithmic scale in Fig. S5a further underscores that our model scales robustly across datasets of varying sizes.

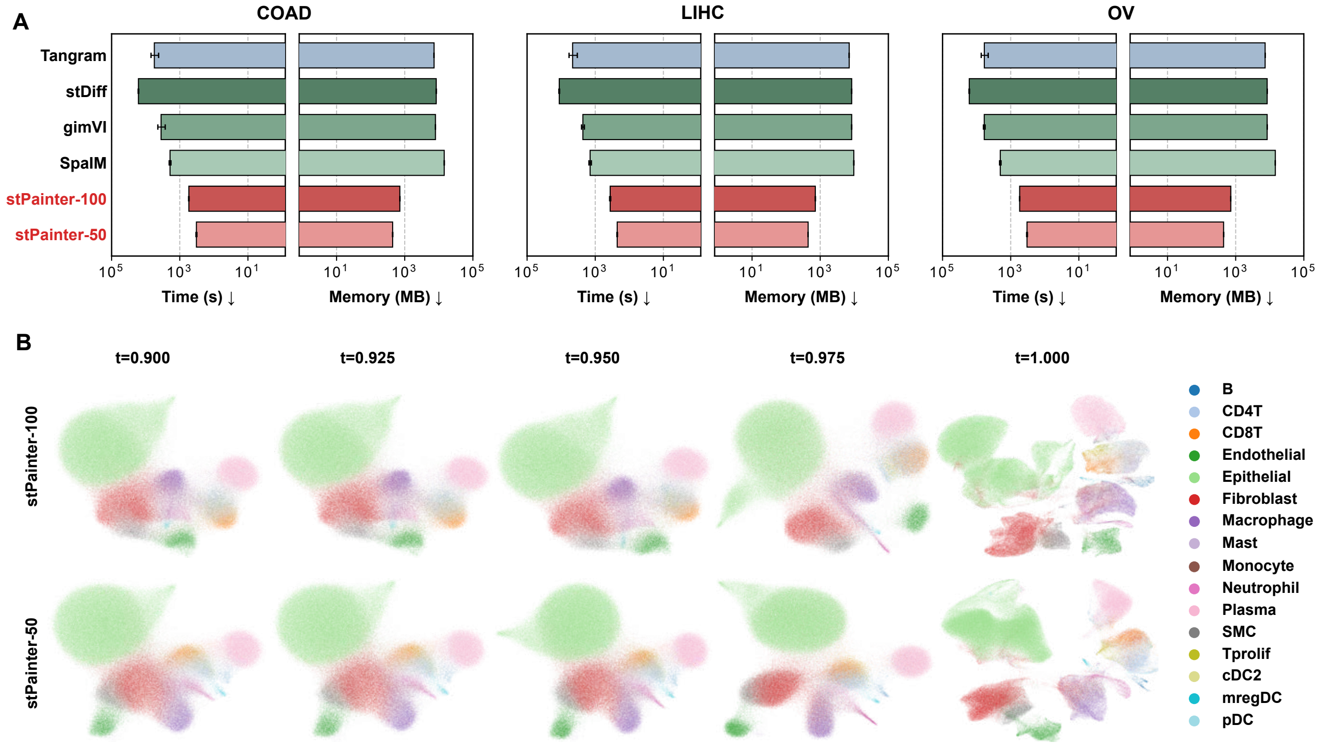


**Fig. S5. Computational efficiency benchmarking and denoising dynamics visualization. a**, Quantitative benchmarking of computational resources across COAD, LIHC, and OV datasets. The bar charts compare the running time (left, seconds) and peak memory usage (right, MB) of STPAINTER variants (STPAINTER-100, STPAINTER-50) against baseline methods. Note that axes are on a logarithmic scale. **b**, UMAP visualization of the progressive denoising trajectory. The sequence illustrates the evolution of latent cellular representations starting from the perturbed state at $t = 0.900$ and gradually refining towards the fully reconstructed manifold at $t = 1.000$. As the process advances, the latent embeddings resolve from unstructured noise into distinct, biologically coherent cell clusters.

We further visualized the generative dynamics of STPAINTER to elucidate how the model reconstructs biological fidelity from noise. Fig. S5b displays the UMAP projection of the latent representations at distinct time steps of the reverse-time SDE process, ranging from the highly perturbed state ($t = 0.900$) to the final target distribution ($t = 1.000$). At the initial stage ($t = 0.900$), the cellular representations appear as unstructured, overlapping distributions, indicating a high degree of entropy. As the denoising process advances (from $t = 0.925$ to $t = 0.975$), the model progressively injects structural information based on the learned pan-cancer priors, causing the latent embeddings to disentangle. By the final time step ($t = 1.000$), the trajectory converges onto a fully reconstructed manifold where distinct, biologically coherent cell clusters emerge. This evolution confirms that STPAINTER effectively learns to reverse the diffusion process, accurately guiding the noisy latent states toward the underlying topology of real cellular populations.

#### S5 Generalizability and Biological Fidelity of STPAINTER Across Cancer Types

To evaluate the generalizability of STPAINTER across diverse tissue architectures, we extended our validation to OV and LIHC datasets. A critical determinant of downstream imputation efficacy is the intrinsic quality of the raw ST data. We first characterized the structural disparities between Xenium-based ST profiles and paired scRNA-seq data by quantifying gene-specific dropout rates.

Across all three cohorts (COAD, LIHC, and OV), we observed a distinct bias: immune-related genes exhibited severe dropout in ST compared to scRNA-seq (e.g., *CD8B* exceeding 99% dropout), whereas epithelial markers showed robust detection in ST (Fig. S6). This discrepancy likely stems from the enzymatic dissociation required for scRNA-seq, which often compromises epithelial cell integrity and transcript recovery. Conversely, the high density of epithelial transcripts in ST can lead to signal spillover, obscuring lower-abundance populations such as immune subsets and complicating unsupervised clustering. While this typically necessitates scRNA-seq-guided transfer annotation, we aimed to demonstrate that data enhancement via STPAINTER enables independent, scRNA-seq-like analytical workflows directly on spatial data without external references.

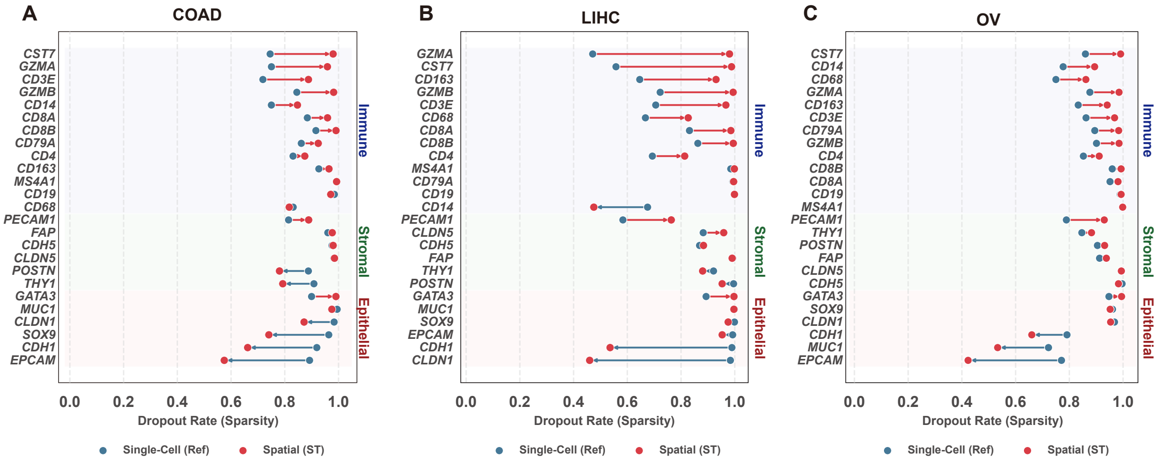


**Fig. S6. The drop-out rates comparison between Xenium dataset and paired scRNA-seq.** The drop-out ratio comparison of COAD (**a**), LIHC (**b**), and OV (**c**).

Following the application of STPAINTER to the OV dataset, we assessed its capacity to resolve cellular heterogeneity. The STPAINTER-enhanced data maintained robust clustering performance for major lineages, particularly when utilizing the learned latent representations (Fig. S7a). The integration of gene imputation with latent embedding facilitated the distinct recovery of lineage-specific marker expression (Fig. S7c). Despite the inherent noise in raw Xenium matrices, which typically results in the commingling of epithelial and non-epithelial signals, STPAINTER significantly mitigated these artifacts. Although some residual overlap persisted in principal component space due to the dense cellular packing of the tumor core (Fig. S7b, d), the model substantially improved major cell type boundary definition on UMAP embeddings. Notably, the inferred TME composition achieved a level of granularity with CODEX dataset comparable to transfer-learning approaches (Fig. S7e), suggesting that STPAINTER retains spatial context even when raw signals are sparse.

At the sub-cluster level, dissecting immune subsets (T cells and myeloid cells) remained challenging due to low capture rates; however, the enhanced data successfully resolved key functional markers, such as *RORC* and *IL26* for Th17 cells, and *HAVCR2* and *LAG3* for exhausted $CD8^+$ T cells, enabling accurate subtype annotation. Conversely, epithelial sub-clustering was precise, demonstrating distinct gene expression gradients and marker specificity. These findings confirm that STPAINTER supports robust cell-type identification even in high-noise spatial environments.

Similarly, within the LIHC landscape (Fig. S9a), the model successfully reconstructed a comprehensive ecosystem comprising hepatocytes, endothelial cells, and diverse immune populations. Crucially, the high-resolution inference allowed for the dissection of the hepatocellular compartment. We distinguished between pericentral and periportal hepatocytes based on metabolic markers, and identified distinct HCC subclones, including proliferative populations marked by *MKI67* (Fig. S10i). This spatial metabolic zonation is often lost in dissociated protocols but was faithfully preserved and enhanced by our model.

Furthermore, the model resolved critical myeloid heterogeneity, effectively separating tissue-resident Kupffer cells from infiltrating macrophages based on canonical markers such as *CD163* and *MARCO*(Fig. S10f). The ability to distinguish these populations is clinically relevant, as their spatial distribution often correlates with prognosis and response to therapy. These results underscore the capacity of STPAINTER to generalize across cancer types while preserving the intricate biological granularity essential for downstream mechanistic interrogation.

To further delineate the functional heterogeneity of epithelial subpopulations, we performed downstream pathway enrichment analyses with a specific focus on epithelial lineage–associated programs. As shown in Supplementary Fig. S11, distinct epithelial subtypes exhibited marked differences in signaling activity, cell-cycle status, metabolic reprogramming, and differentiation states. Stem-like and proliferative epithelial subsets were characterized by robust activation of WNT/$\beta$-catenin signaling pathways, accompanied by elevated cell-cycle and proliferation scores. Differentiated epithelial populations displayed higher enrichment of differentiation-associated and metabolic pathways, such as gluconeogenesis and lipid metabolism programs in LIHC dataset. Invasive or mesenchymal-like epithelial cells showed pronounced activation of epithelial–mesenchymal transition (EMT) and TGF-$\beta$ signaling, alongside hypoxia-responsive and glycolytic pathways. Collectively, these pathway-level distinctions underscore the functional diversification among epithelial subpopulations and provide independent support for the downstream analyses of imputed genesets following stPainter enhancement.

Downstream analyses of datasets processed by alternative imputation tools revealed fundamental structural deviations from standard single-cell resolution ST matrices. Specifically, artificial data densification led to near-complete loss of drop-out structure, wherein 100% of genes exhibid erroneously assigned SMC markers (*RGS5*, *NOTCH3*) to endothelial cells, whereas Tangram resulted in pervasive leakage of epithelial signatures (*KRT18*, *EPCAM*) into non-epithelial subsets, often surpassing the expression intensities observed in actual epithelial cells (Fig. S12c). Collectively, these findingted expression (Fig. S12a-c). Unlike stPainter, which explicitly models the inherent drop-out characteristics of ST data, most existing methods did not preserve this feature. Moreover, we identified some distortions in lineage-specific marker fidelity. In the LIHC dataset, gimVI yielded imputed expression of B-cell markers (*CD19*, *CD79A*) in proliferating T cells and cDC1 at levels exceeding those in B cells and plasma cells, while Tangram exhibited ectopic expression of *FAP* in endothelial cells and *KRT18* in SMCs (Fig. S12a). Similarly, for the OV dataset, gimVI attenuated Mast cell signatures while inducing indiscriminate co-expression of SMC and fibroblast markers (*PDGFRA*, *FAP*, *PDGFRB*) within the stromal compartment; Tangram exhibited confounding lineage mixing characterized by the co-expression of *CD7* and *CD34* in both vascular endothelial cells and $CD4^+$ T cells (Fig. S12b). Furthermore, in the COAD dataset, gimVI failed to recover T-cell markers (*CD7*, *CD3E*, *CD3D*) ans indicate that current alternative tools compromise downstream analytical utility; in contrast, stPainter generates single-cell resolution ST data that faithfully recapitulates native biological characteristics and pipeline compatibility, highlighting the advantages of stPainter in preserving biologically meaningful structure for spatial analysis.

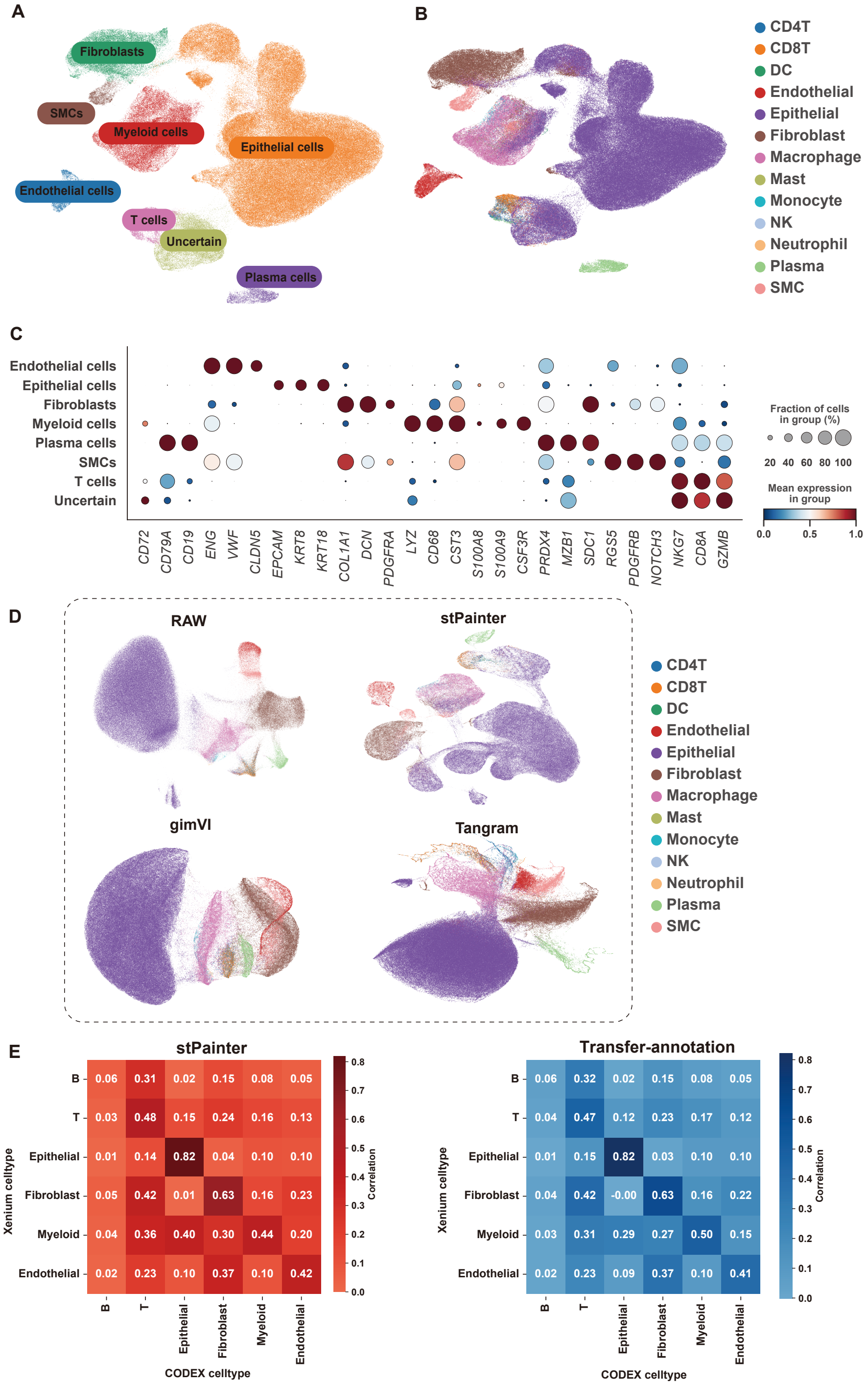


**Fig. S7. Major celltypes identification of the Ovarian Cancer Xenium dataset.** **a**, UMAP embedding of annotated major cell lineages in OV xenium dataset. **b**, UMAP embedding of transfer-annotation cell lineages in OV xenium dataset. **c**, Dotplot demonstrating the expression of gene markers of major celltypes. **d**,Comparative UMAP topologies generated by different computational workflows (STPAINTER-100, gimVI, and Tangram). **e**,The heatmap of cell-type proportions and correlation matrices (Pearson's *r* index) validate the concordance between STPAINTER-imputed cellular compositions and ground-truth CODEX proteomics across tissue patches.

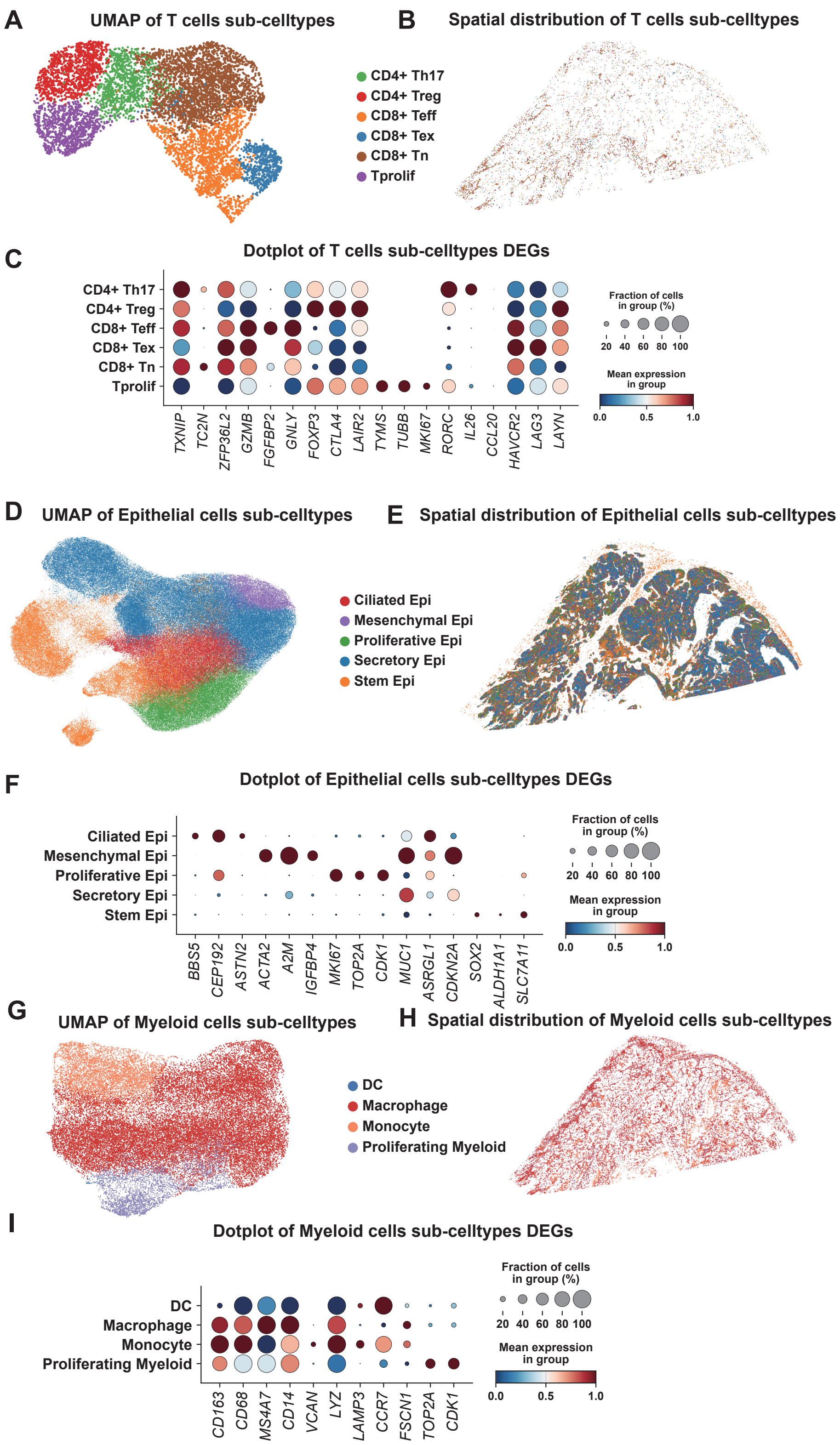


**Fig. S8. Fine-grained dissection of cellular heterogeneity in Ovarian Cancer. a-c**, UMAP projection, spatial distribution and DEGs of T cell subtypes. **d-f**, UMAP projection, spatial distribution and DEGs of Epithelial cell subtypes. **g-i**, UMAP projection, spatial distribution and DEGs of Myeloid cell subtypes.

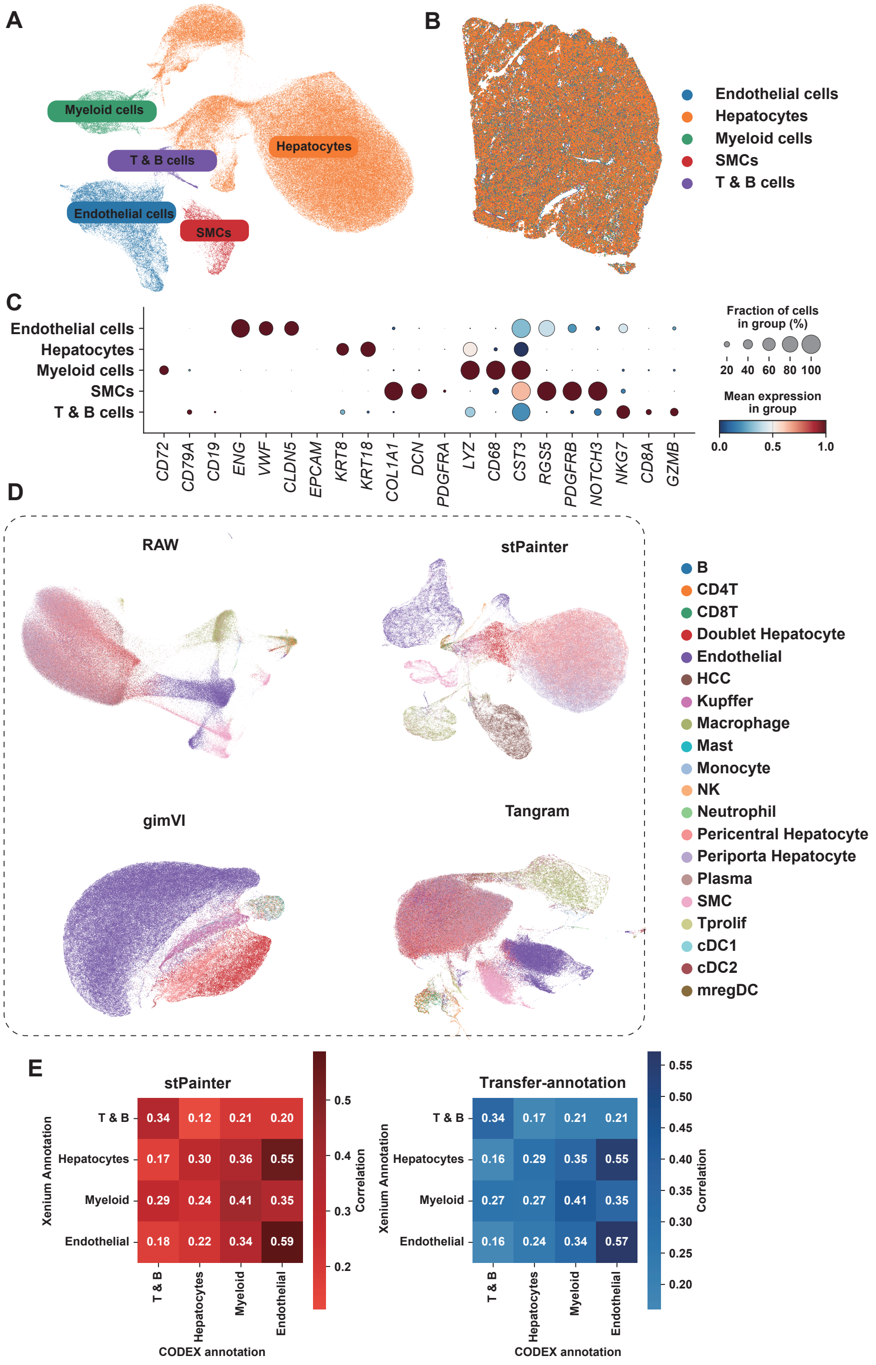


**Fig. S9. Major celltypes identification of the Liver Hepatocellular Carcinoma Xenium dataset.** **a**, UMAP embedding of annotated major cell lineages in LIHC xenium dataset. **b**, UMAP embedding of transfer-annotation cell lineages in LIHC xenium dataset. **c**, Dotplot demonstrating the expression of gene markers of major celltypes. **d**, Comparative UMAP topologies generated by different computational workflows (STPAINTER-100, gimVI, and Tangram). **e**, The heatmap of cell-type proportions and correlation matrices (Pearson's $r$ index) validate the concordance between STPAINTER-imputed cellular compositions and ground-truth CODEX proteomics across tissue patches.

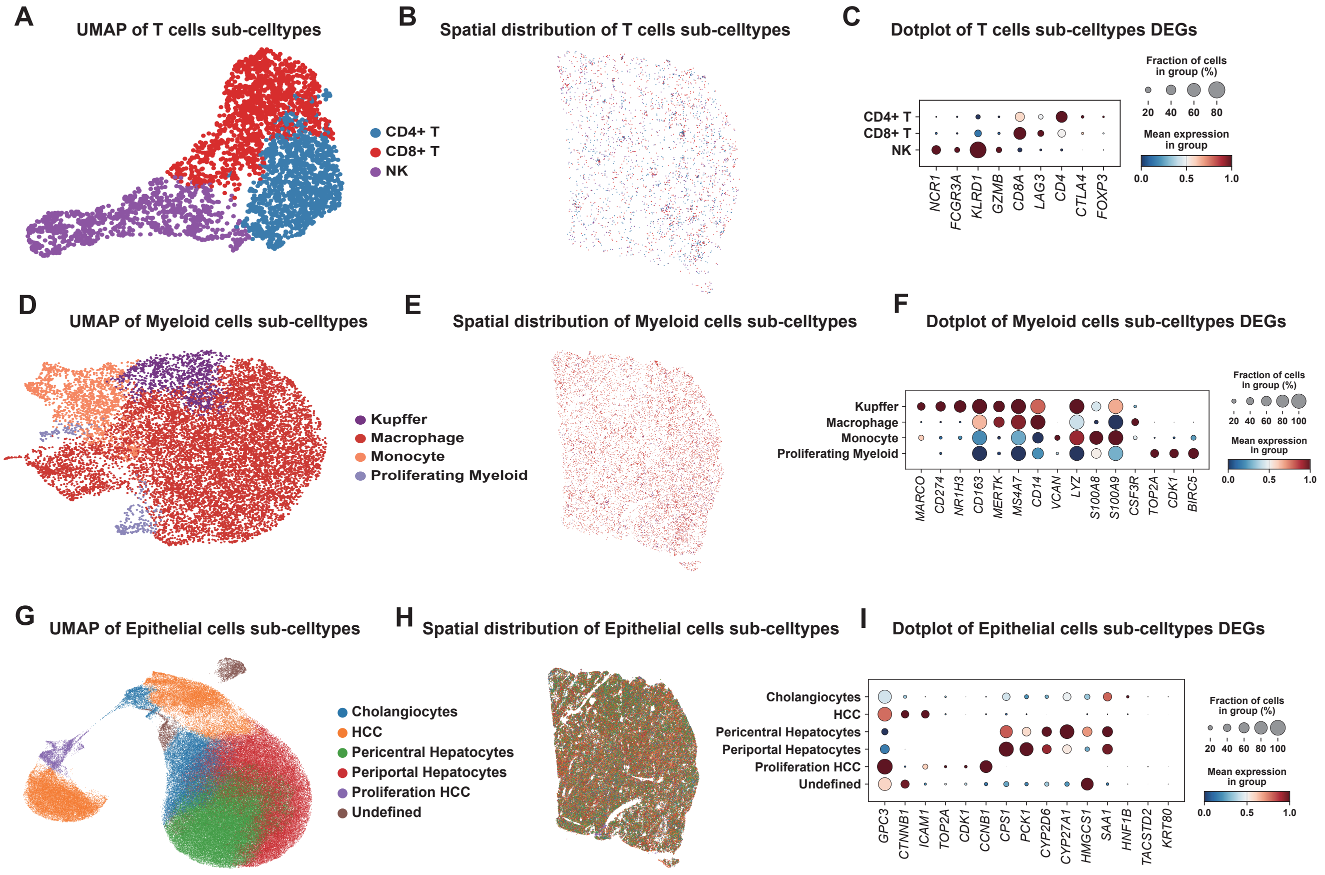


**Fig. S10. Fine-grained dissection of cellular heterogeneity in LIHC. a-c**, UMAP projection, spatial distribution and DEGs of T cell subtypes. **d-f**, UMAP projection, spatial distribution and DEGs of Myeloid cell subtypes. **g-i**, UMAP projection, spatial distribution and DEGs of Epithelial cell subtypes.

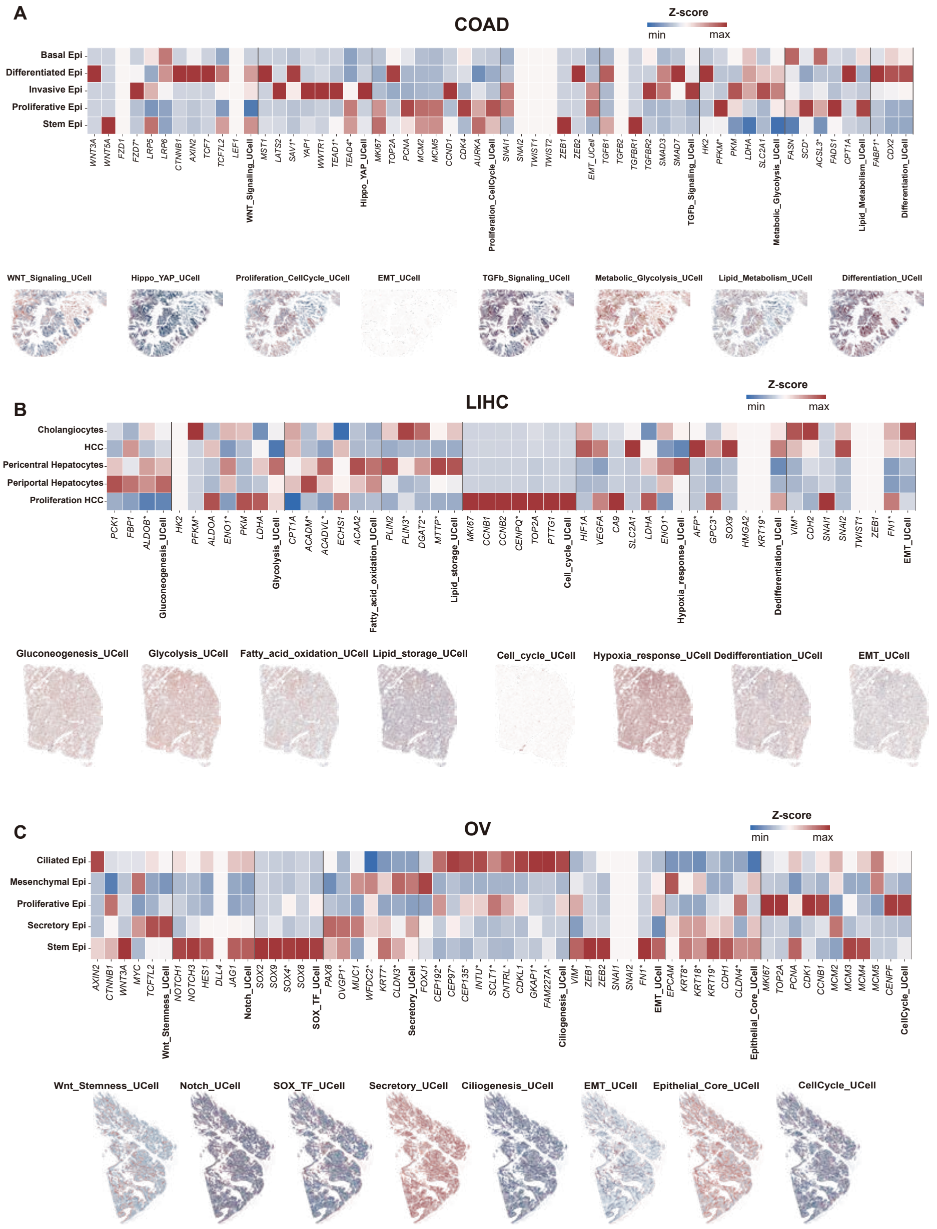


**Fig. S11. Pathway enrichment analysis of epithelial cell subtypes. a**, Heatmap showing enriched pathways and associated genes across epithelial subtypes in the COAD Xenium dataset, together with the spatial distribution of pathway enrichment scores. *indicates imputed genes. **b**, Heatmap showing enriched pathways and associated genes across epithelial subtypes in the LIHC Xenium dataset, together with the spatial distribution of pathway enrichment scores. *indicates imputed genes. **c**, Heatmap showing enriched pathways and associated genes across epithelial subtypes in the OV Xenium dataset, together with the spatial distribution of pathway enrichment scores. *indicates imputed genes.

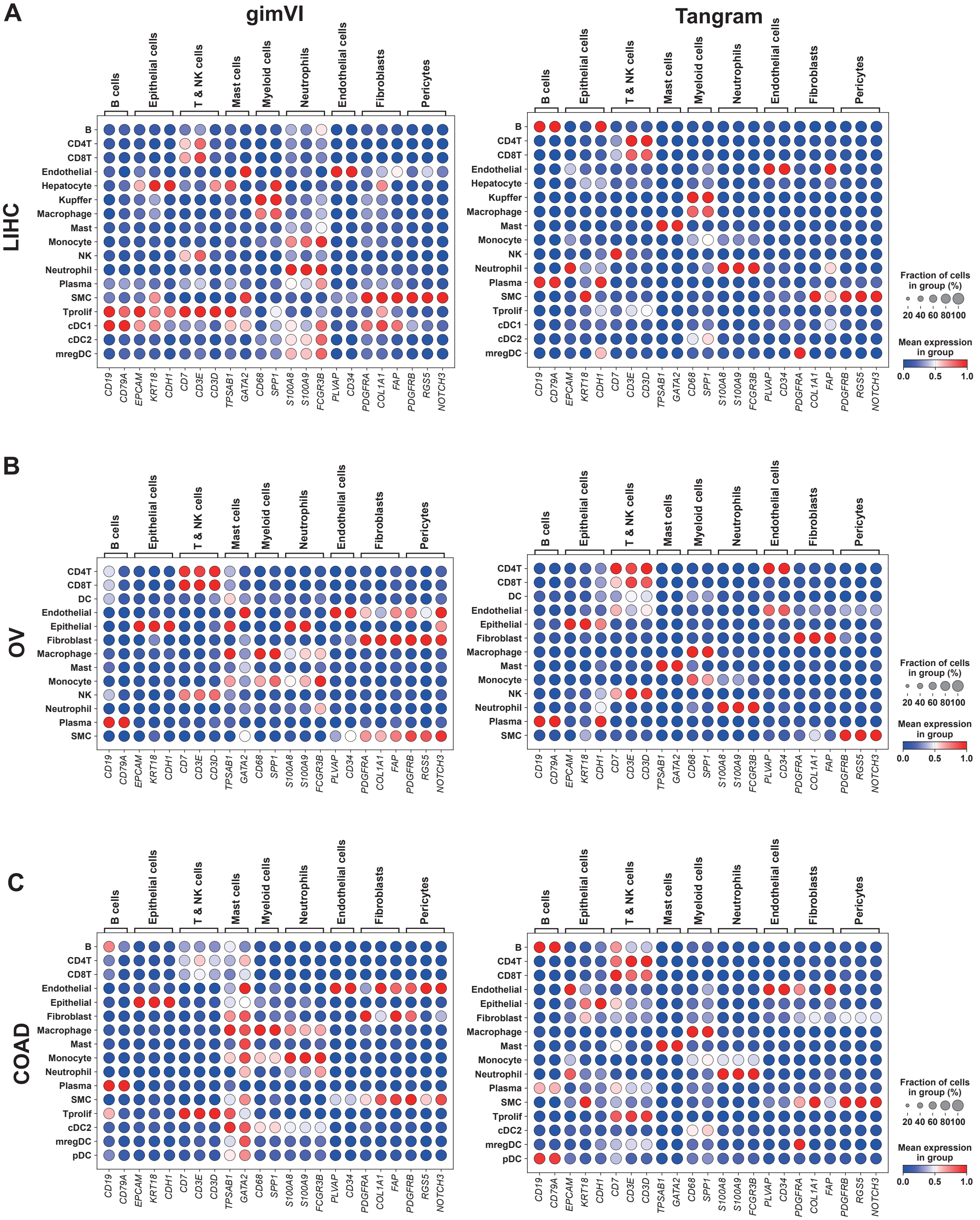


**Fig. S12. Fine-grained dissection of cellular heterogeneity in LIHC. a**, dotplots of DEGs of major celltypes in LIHC dataset imputed by gimVI and Tangram. **b**, dotplots of DEGs of major celltypes in OV dataset imputed by gimVI and Tangram. **c**, dotplots of DEGs of major celltypes in COAD dataset imputed by gimVI and Tangram.
